# Intron-assisted, viroid-based production of insecticidal circular double-stranded RNA in *Escherichia coli*

**DOI:** 10.1101/2020.12.02.407684

**Authors:** Beltrán Ortolá, Teresa Cordero, Xu Hu, José-Antonio Daròs

## Abstract

RNA interference (RNAi) is a natural mechanism for protecting against harmful genetic elements and regulating gene expression, which can be artificially triggered by the delivery of homologous double-stranded RNA (dsRNA). This mechanism can be exploited as a highly specific and environmentally friendly pest control strategy. To this aim, systems for producing large amounts of recombinant dsRNA are necessary. We describe a system to efficiently produce large amounts of circular dsRNA in *Escherichia coli* and demonstrate the efficient insecticidal activity of these molecules against Western corn rootworm (WCR, *Diabrotica virgifera virgifera* LeConte), a highly damaging pest of corn crops. In our system, the two strands of the dsRNA are expressed in *E. coli* embedded within the very stable scaffold of *Eggplant latent viroid* (ELVd), a small circular non-coding RNA. Stability in *E. coli* of the corresponding plasmids with long inverted repeats was achieved by using a cDNA coding for a group-I autocatalytic intron from *Tetrahymena thermophila* as a spacer. RNA circularization and large-scale accumulation in *E. coli* cells was facilitated by co-expression of eggplant tRNA ligase, the enzyme that ligates ELVd during replication in the host plant. The inserted intron efficiently self-spliced from the RNA product during transcription. Circular RNAs containing a dsRNA moiety homologous to *smooth septate junction 1* (*DvSSJ1*) gene exhibited excellent insecticide activity against WCR larvae. Finally, we show that the viroid scaffold can be separated from the final circular dsRNA product using a second *T. thermophila* self-splicing intron in a permuted form.

## Introduction

RNA silencing, also known as RNA interference (RNAi), is a eukaryotic natural defense mechanism against exogenous RNA and transposon mobilization that has evolved to also regulate gene expression. RNAi is induced by the presence of highly structured or double-stranded RNA (dsRNA) and typically results in the silencing of homologous genes [1]. Since efficient silencing can be equally induced by endogenously transcribed or exogenously delivered RNA, RNAi-mediated gene knockdown is frequently used in many organisms for basic research to study gene function, as well as for biotechnological applications, from therapeutics [2] to plant breeding [3]. More specifically, in recent years, remarkable progress has been made in the use of exogenously supplied dsRNA as a highly specific and environmentally friendly anti-pest and anti-pathogen agent in agriculture [4–6]. The ingestion of long dsRNAs by nematodes, insects, or other arthropods induces silencing of endogenous homologous genes, which may cause pest death or, at least, affect development, feeding, mobility, or progeny production, reducing crop damage in any case [7].

The dsRNA molecules required for RNAi applications can be obtained via chemical synthesis or bi-directional *in vitro* transcription. Both strategies generate two complementary RNAs that must be subsequently hybridized. These strategies are time-consuming, expensive, and particularly difficult to scale up to produce the large amounts of dsRNAs required, for example, in pest control. A more feasible strategy is *in vivo* production using a biofactory system, such as the bacteria *Escherichia coli* [8]. In this approach, the dsRNA can be expressed from a single transcriptional unit, which results in a hairpin RNA consisting of two complementary strands of the target sequence separated by a single-stranded loop [9–12]. However, the presence of inverted repeats in plasmid vectors significantly damages stability [13,14]. Alternatively, the two complementary RNA strands are usually synthesized *in vivo* from two promoters in inverted orientations [15–17]. Again, this strategy requires hybridization of both complementary strands, thereby lowering efficiency and rendering the whole process prone to degradation.

We have recently developed a system to produce large amounts of recombinant RNA in *E. coli* based on elements of viroid biology [18,19]. Viroids are a unique class of plant infectious agents that are exclusively composed of a relatively small (246–434 nt) circular non-coding RNA molecule [20–22]. Our RNA production system is based on co-expression in *E. coli* of an *Eggplant latent viroid* (ELVd) [23] scaffold, in which the RNA of interest is grafted, along with the eggplant (*Solanum melongena* L.) tRNA ligase, the host enzyme involved in viroid circularization in the infected plant [24,25]. Although there is no ELVd replication in *E. coli*, the viroid-derived RNA can be efficiently transcribed in these bacteria and it undergoes processing through the embedded hammerhead ribozymes. The resulting monomers that contain the RNA of interest are recognized by the tRNA ligase and circularized. The expression product likely remains bound to the tRNA ligase, forming a ribonucleoprotein complex that reaches high concentration in *E. coli* cells. Using this system, tens of milligrams of RNAs of interest, such as RNA aptamers, can be easily obtained per liter of *E. coli* culture under regular laboratory conditions [18,26]. We aim to apply this system for producing the large amounts of dsRNAs required to fight Western corn rootworm (WCR; *Diabrotica virgifera virgifera* LeConte; Coleoptera: Chrysomelidae), using RNAi strategies. WCR is considered one of the most harmful insect pests of cultivated corn in the USA and it has received increasing attention globally because of repeated invasion events outside this country [27]. However, despite our initial success in producing recombinant hairpin RNAs of small length, we experienced major difficulties in building the expression plasmids that contain long inverted repeats. Thus, we first aimed to adapt the viroid-based system for the large-scale production in *E. coli* of hairpin RNAs with long double-stranded regions, which are required in anti-pest RNAi approaches. Second, we sought to use these recombinant RNAs to fight WCR. Third, we refined the *in vivo* production system to automatically remove the viroid scaffold from the dsRNA product.

We show that plasmids with long inverted repeats become completely stable in *E. coli* when the corresponding sequences are separated by a sequence coding for a group-I self-splicing intron. Interestingly, this intron self-splices with extremely high efficiency after transcription *in vivo*, facilitating the formation of an RNA product that contains the long hairpin RNA and that accumulates to high concentration in *E. coli*. We also demonstrate that the resulting RNA product, which consists of a structured viroid-derived scaffold from which the dsRNA protrudes, shows a potent insecticide activity against WCR larvae. Finally, we show that the dsRNA of interest can be efficiently excised from the viroid scaffold through the addition of a permuted version of the same intron flanking the inverted repeats; this yields a highly-stable and compact circular molecule consisting of a perfect dsRNA locked at both ends with small terminal single-stranded loops.

## Results

### Plasmids with long inverted repeats are stabilized in *E. coli* when these sequences are separated by a cDNA corresponding to a self-splicing group-I intron

*D. virgifera smooth septate junction 1* (*DvSSJ1*) gene encodes a membrane protein associated with smooth septate junctions, which are required for intestinal barrier function. Ingestion by WCR larvae of dsRNA homologous to *DvSSJ1* induces mRNA suppression and larval growth inhibition and mortality [28–30]. Using the viroid-derived system to produce recombinant RNA in *E. coli*, we attempted to produce the large amounts of various dsRNAs homologous to *DvSSJ1* to analyze their anti-WCR activity via oral feeding of insect larvae. However, we were unable to obtain the corresponding expression plasmids with long inverted repeat sequences, although we tried multiple cloning strategies, *E. coli* strains, and growth conditions. We reasoned that plasmid instability would revert if inverted repeats were separated by a sufficiently long spacer sequence. Inspired by previous work to produce hairpin RNAs in plants—in which inverted repeats were separated by a cDNA corresponding to a plant intron that efficiently spliced when the RNA was transcribed in the plant cells [31]—we searched for introns potentially able to self-splice in *E. coli*. We selected the group-I *Tetrahymena thermophila* 26S rRNA intron [32].

In contrast to previous fruitless results, we easily obtained a plasmid in which 83-nt inverted repeats homologous to *DvSSJ1* were separated by a 433-bp cDNA that corresponded to the *T. thermophila* 26S rRNA intron (GenBank accession number V01416.1), plus both 10-nt native flanking exons. To build this plasmid, we electroporated the product of a Gibson assembly reaction in *E. coli* and selected transformed clones in plates containing ampicillin. Electrophoretic analysis of recombinant plasmids from 12 independent *E. coli* colonies showed that all were the same size and exhibited a migration delay consistent with the inserted cDNA (Supplemental Figure S1, compare lane 1 with lanes 2 to 13). The expected sequence was confirmed in one of these plasmids, hereafter named pLELVd-DvSSJ1 (Supplemental Dataset S1).

### Remarkable amounts of the dsRNA of interest, inserted into the ELVd molecule, accumulate in *E. coli* when co-expressed with tRNA ligase

We used pLELVd-DvSSJ1 to co-electroporate the RNase III-deficient strain of *E. coli* HT115(DE3), along with plasmid p15LtRnlSm (Supplemental Dataset S1), from which eggplant tRNA ligase is constitutively expressed. As controls, p15LtRnlSm was also co-electroporated with the empty expression plasmid (pLPP) or the plasmid to express empty ELVd (pLELVd), with no RNA of interest inserted (Supplemental Dataset S1). Three independent colonies were selected from plates containing ampicillin and chloramphenicol and grown for 24 h in Terrific Broth (TB). We extracted total RNA from the cells and analyzed it by polyacrylamide gel electrophoresis (PAGE) in denaturing conditions (8 M urea). Two prominent bands above the 600-nt and slightly below the 400-nt RNA markers were observed in the lanes containing RNA from bacteria transformed with pLELVd-DvSSJ1 (Figure 1(a), lanes 7 to 9, orange and black arrows, respectively). Note that these bands exhibited a fluorescence signal higher than those corresponding to endogenous *E. coli* rRNAs (Figure 1(a), upper part of the gel), indicating a large accumulation *in vivo*. RNA extracts from the empty ELVd controls exhibited a single prominent band above the 400-nt marker that, according to our previous analyses [33,34], corresponds to the 333-nt circular ELVd RNA (Figure 1(a), lanes 4 to 6, white arrow). In denaturing conditions, circular RNAs migrate more slowly than the linear counterparts of the same size. Finally, RNA extracts from the empty plasmid control did not exhibit any particular prominent band (Figure 1(a), lanes 1 to 3).

**Figure 1.**
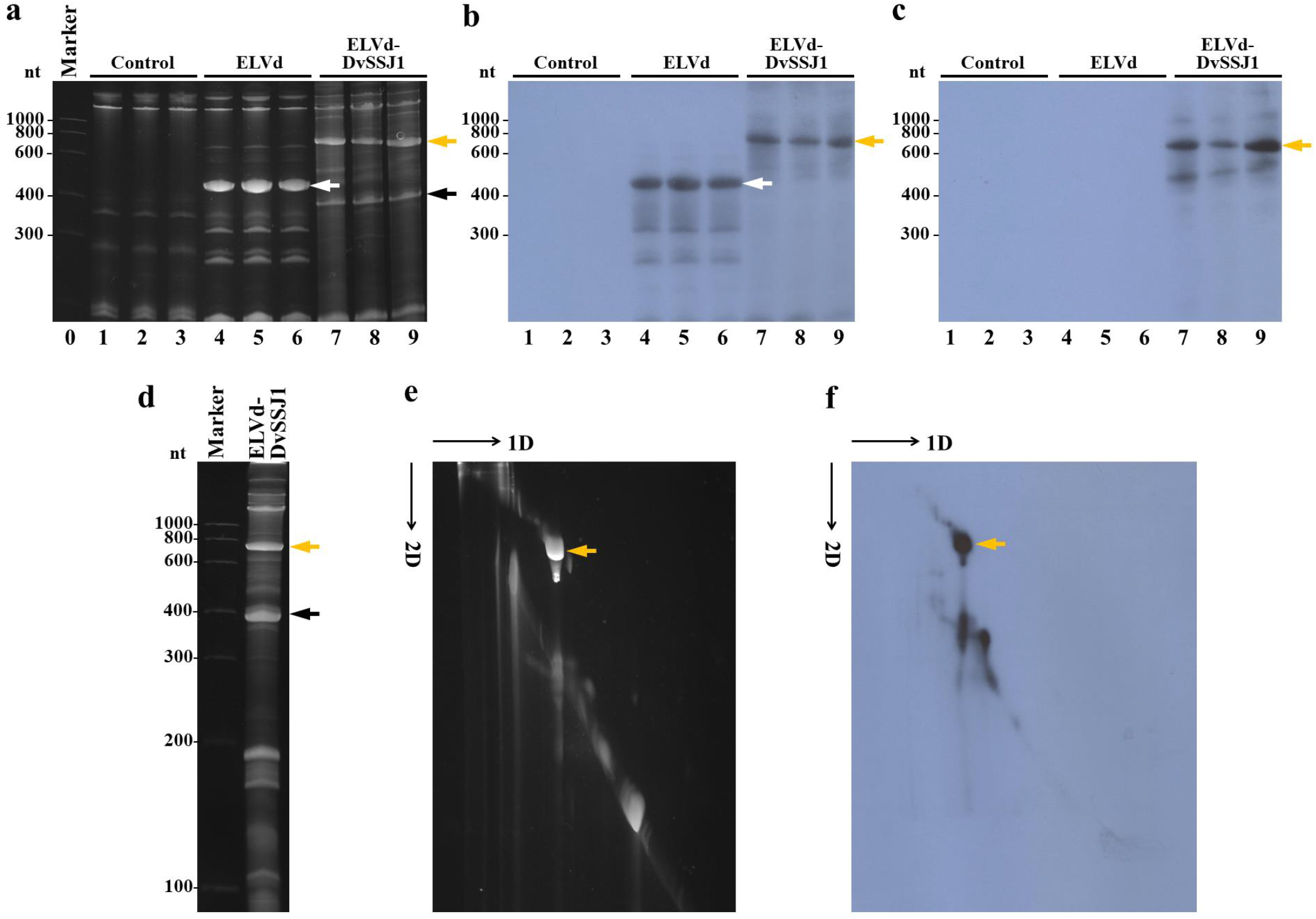
Analysis of *DvSSJ1*-derived dsRNA produced in *E. coli*. (a to c) Bacterial RNA was extracted from three independent cultures of *E. coli* co-transformed with p15LtRnlSm and pLPP (lanes 1 to 3), pLELVd (lanes 4 to 6) or pLELVd-DvSSJ1 (lanes 7 to 9), separated by denaturing PAGE, and stained with ethidium bromide (a) or transferred to membranes for hybridization with ELVd (b) or *DvSSJ1* (c) probes. (a) Lane 0, RNA marker with sizes in nt on the left. (d to f) An RNA preparation from *E. coli* co-transformed with p15LtRnlSm and pLELVd-DvSSJ1 was separated by denaturing 2D PAGE. The first dimension was under high ionic strength, and the RNAs were stained with ethidium bromide (d). The second dimension was under low ionic strength; the RNAs were first stained with ethidium bromide (e) and then transferred to a membrane and hybridized with a *DvSSJ1* probe (f). (e and f) Slim black arrows indicate the direction of RNA migration in both dimensions of 2D PAGE. Thick white, orange, and black arrows point to ELVd, ELVd-DvSSJ1, and spliced intron, respectively.

To confirm the identity of the expressed RNA species, we also analyzed the RNA preparations by northern blot hybridization, using radioactive RNA probes complementary to ELVd and the sense strand of the *DvSSJ1*-derived dsRNA. While the ELVd probe hybridized with the prominent RNA band in the empty ELVd controls and the slowly migrating band in the ELVd-DvSSJ1 samples (Figure 1(b), lanes 4 to 9, white and orange arrows, respectively), the DvSSJ1 probe only hybridized with the slowly migrating band of the ELVd-DvSSJ1 samples (Figure 1(c), lanes 7 to 9, orange arrow). These results indicate that this slowly migrating band corresponds to a composite RNA species consisting of ELVd and *DvSSJ1* moieties. To determine whether this RNA species was linear or circular, we separated an RNA preparation from bacteria co-transformed with p15LtRnlSm and pLELVd-DvSSJ1 using denaturing 2-dimension (2D) PAGE, under conditions of high ionic strength (Figure 1(d)) and then low ionic strength. In this electrophoretic separation, circular RNAs are selectively delayed when conditions change from high to low ionic strength; they deviate from the diagonal of the linear RNAs. We observed the electrophoretic behavior of a circular RNA for the prominent slowly migrating species when, after the second run, the gel was either stained with ethidium bromide (Figure 1(e), orange arrow) or hybridized with the *DvSSJ1* probe (Figure 1(f), orange arrow). Hybridization spots in the diagonal of linear molecules must correspond to linear counterparts of the ELVd-DvSSJ1 RNA circular form (Figure 1(f)).

We also sought to determine the nature of the rapidly migrating prominent band in the ELVd-DvSSJ1 samples (Figure 1(a), lanes 7 to 9, black arrow). An RNA preparation from *E. coli* co-transformed with pLtRnlSm and pLELVd-DvSSJ1 was separated via denaturing PAGE (Figure 2(a)) and hybridized with a probe complementary to the *T. thermophila* 26S rRNA group-I intron. The probe specifically recognized this rapidly migrating species in the ELVd-DvSSJ1 RNA preparation (Figure 2(b), black arrow), indicating that this band corresponds to the spliced intron that also accumulates to a high concentration in *E. coli*. The electrophoretic mobility of this species (close to the 400-nt RNA marker) suggests a linear form. The full size of the spliced *T. thermophila* 26S rRNA intron is 413 nt. We were surprised that we did not obtain hybridization bands corresponding to unspliced forms of the intron. This suggests an extremely efficient self-splicing reaction in *E. coli* cells. To investigate the efficiency of intron splicing *in vivo*, we sampled a liquid *E. coli* culture at several time points and analyzed bacterial RNA by electrophoretic separation and northern blot hybridization with the *DvSSJ1* and the intron probes (Figure 2(c)). Time-course analysis definitively supported a highly efficient self-splicing reaction of the *T. thermophila* intron in *E. coli*. No substantial amounts of processing intermediates were detected at any time point. A deletion analysis of the flanking 10-nt exon fragments suggested that their size could also be reduced with no substantial effect on self-cleavage (Supplemental Figure S2).

**Figure 2.**
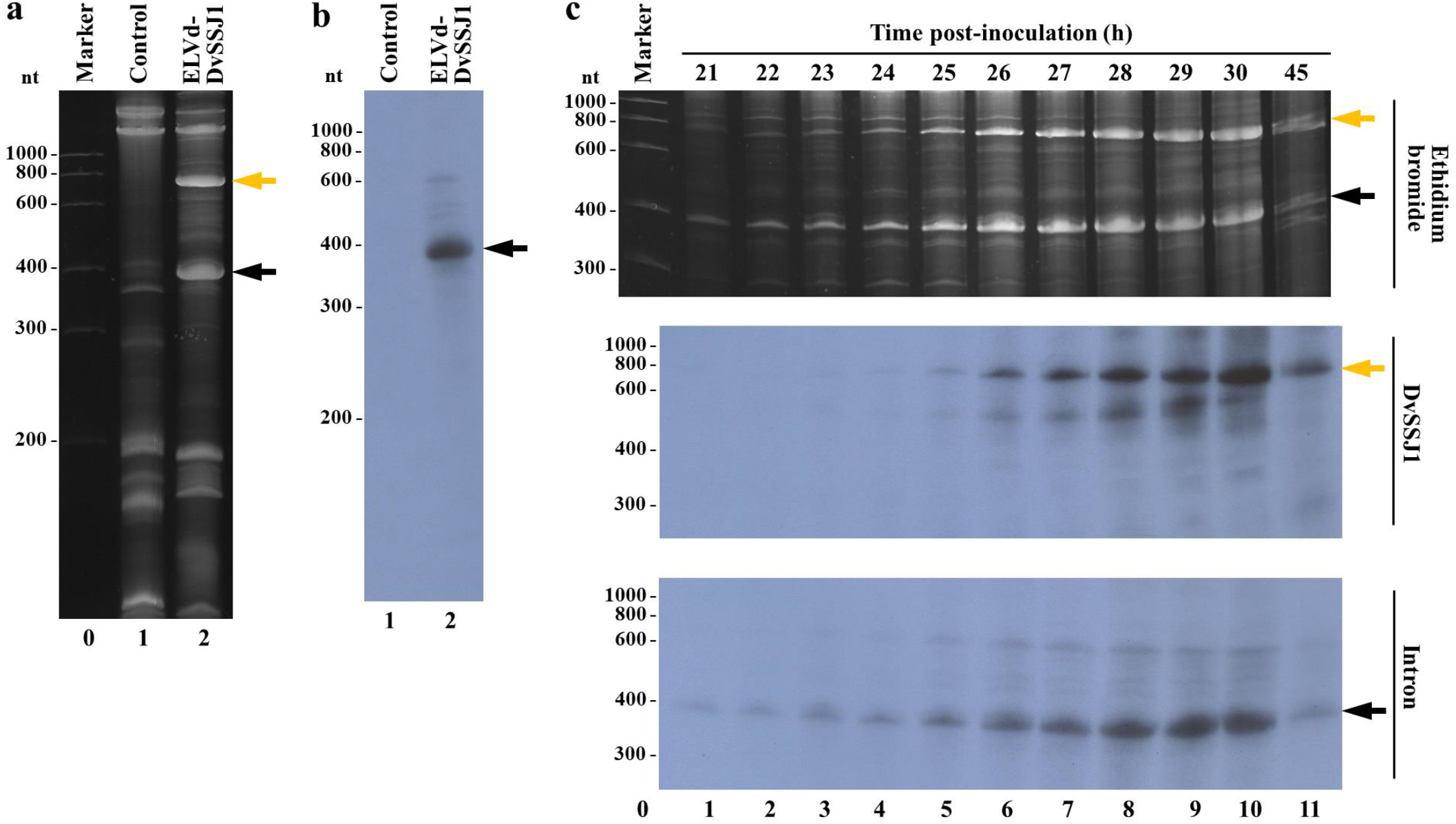
Analysis of the 26S rRNA intron processing in *E. coli*. (a and b) RNA was extracted from *E. coli* transformed with p15LtRnlSm and either pLPP (lane 1) or pLELVd-DvSSJ1 (lane 2), separated by denaturing PAGE, stained with ethidium bromide (a), and then transferred to a membrane and hybridized with an intron-specific probe (b). (c) RNA was extracted from aliquots of a liquid culture of *E. coli* co-transformed with p15LtRnlSm and pLELVd-DvSSJ1 at different time points (as indicated). Lanes 1 to 11, RNAs were separated by denaturing PAGE and stained with ethidium bromide (top) or transferred to a membrane and hybridized with a *DvSSJ1* (middle) or intron-specific (bottom) probe. Orange and black arrows point to ELVd-DvSSJ1 RNA and the spliced intron, respectively. (a and c) Lane 0, RNA markers with sizes in nt on the left.

These results indicate that, while the *T. thermophila* cDNA serves to stabilize the inverted repeats in the *E. coli* expression plasmids, the intron very efficiently self-splices from the primary transcript in bacterial cells, facilitating the accumulation of a circular RNA product that consists of an ELVd scaffold from which the *DvSSJ1*-derived 83-bp hairpin RNA is presented. The entire process is schematized in Figure 3.

**Figure 3.**
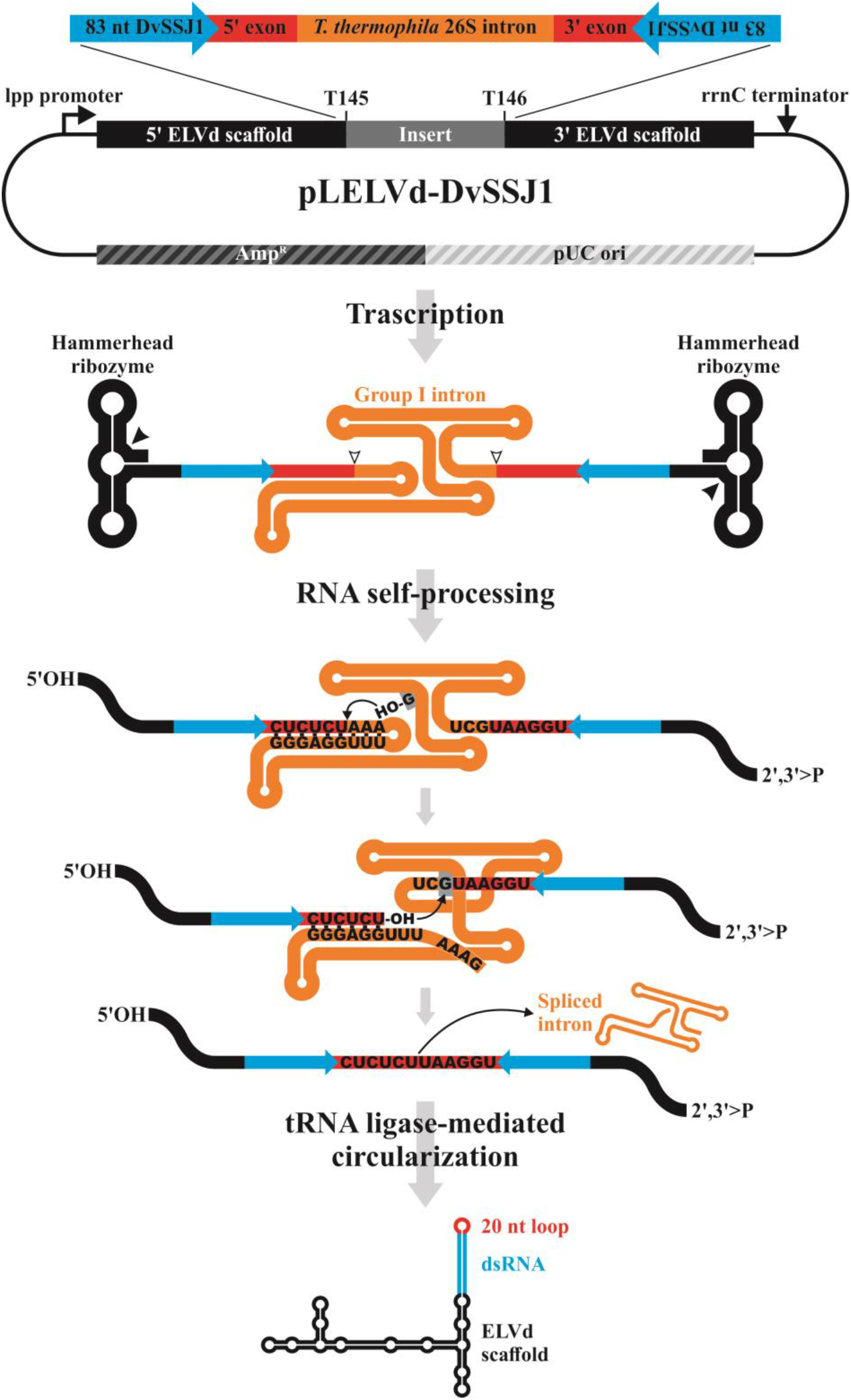
Schematic representation of the pLELVd-DvSSJ1 plasmid and the process for producing dsRNA in *E. coli*. In the primary transcript (not at scale), the inverted repeats homologous to *DvSSJ1* are separated by the *T. thermophila* 26S rRNA intron and the 10-nt native flanking exons. Spacing the inverted repeats in the expression plasmid is critical to stabilizing the constructs. After transcription, the intron self-splices very efficiently through two sequential transesterification reactions. First, the 3’-OH of an exogenous guanosine nucleoside docked in the intron structure attacks the phosphodiester bond between the first exon-intron boundary, generating a 3’-OH group at the end of the exon and leaving the G residue attached to the 5’ end of the intron. Next, the intron terminal G is docked in the same place of the intron structure and the 3’-OH group of the first exon attacks the phosphodiester bond between the second exon-intron boundary, resulting in the ligation of both exons and the release of the catalytic intron in a linear form. The intron-processing facilitates the hybridization of the inverted repeat sequences to form a hairpin composed of a dsRNA capped by a 20-nt loop arising from the two flanking exons. Concomitantly, the self-splicing activity of the two ELVd hammerhead ribozymes present in the precursor yields the 5’-hydroxyl and 2’,3’-phosphodiester termini that are recognized and ligated by the co-expressed eggplant tRNA ligase, generating a circular chimera.

### The ELVd-DvSSJ1 RNA produced in *E. coli* possesses insecticide activity against WCR larvae

To test whether the chimeric ELVd-DvSSJ1 RNA displays anti-WCR activity, we performed a bioassay with WCR larvae. First, we grew 250-ml cultures of *E. coli* co-transformed with p15LtRnlSm and either pLELVd or pLELVd-DvSSJ1. The cells were harvested at 24 h and total bacterial RNA was purified. Our electrophoresis dilution analysis, along with a comparison to standards of known concentration, allowed us to quantify the concentration of empty ELVd and ELVd-DvSSJ1 in both RNA preparations (Supplemental Figure S3). As a control for this assay, we also produced the same *DvSSJ1*-derived 83-bp dsRNA that is contained in ELVd-DvSSJ1, using conventional *in vitro* transcription and hybridization [28]. Next, equivalent amounts of the three RNA preparations were mixed with the artificial rootworm diet. While empty ELVd had no effect on larval growth at the top dose of 35 ng/μl, ELVd-DvSSJ1 and *in vitro*-transcribed *DvSSJ1* induced similar larval growth inhibition (50% inhibition concentration, IC_50_ = 0.159 vs. 0.215 ng/μl) and mortality (50% lethal concentration, LC_50_ = 0.642 vs. 0.665 ng/μl) (Table 1).

**Table 1.** Insecticidal activity of conventional *DvSSJ1 in vitro*-transcribed dsRNA (IVT DvSSJ1) and ELVd-DvSSJ1 dsRNA against WCR

| WCR | LC<sub>50</sub>/IC<sub>50</sub> * | RNA (ng/μl) | Lower 95% CL * | Upper 95% CL | n |
| --- | --- | --- | --- | --- | --- |
| **IVT DvSSJ1** | LC<sub>50</sub> | 0.665 | 0.426 | 0.928 | 188 |
|  | IC<sub>50</sub> | 0.215 | 0.096 | 0.301 | 157 |
| **ELVd-DvSSJ1** | LC<sub>50</sub> | 0.642 | 0.384 | 0.925 | 188 |
|  | IC<sub>50</sub> | 0.159 | 0.028 | 0.277 | 124 |
| **Empty ELVd** | LC<sub>50</sub> | >35 ppm |  |  |  |
|  | IC<sub>50</sub> | >35 ppm |  |  |  |

\*LC<sub>50</sub>, 50% lethal concentration; IC<sub>50</sub>, 50% inhibition concentration; CL, confidence limit.

### Production of a circular version of the dsRNA of interest without the viroid scaffold using a permuted intron

Because carrying a sequence derived from an infectious agent may not be desirable for the commercial use of recombinant RNAs, we next aimed to automatically remove, *in vivo*, the viroid scaffold moiety from the dsRNA product. For this purpose, we inserted into our construct an additional copy of the *T. thermophila* group-I autocatalytic intron, albeit with permuted intron-exon (PIE) sequences [35]. More specifically, we incorporated cDNAs corresponding to the 3’ half of the *T. thermophila* intron (from position C236 in the intron to C10 of exon 2) just between the end of the 5’ ELVd moiety and the sense copy of the inverted repeat; between the antisense copy of the inverted repeat and the start of the 3’ ELVd moiety, we inserted the 5’ half of the *T. thermophila* intron (from −10A of exon 1 up to intron U235), as depicted in Figure 4. The two intron halves were still able to recognize the intron-exon boundaries and undergo the two transesterification reactions. Because the 3’ end of the second exon is covalently linked to the 5’ end of the first exon, both exons and the sequence between them are released as a circular RNA molecule [35].

**Figure 4.**
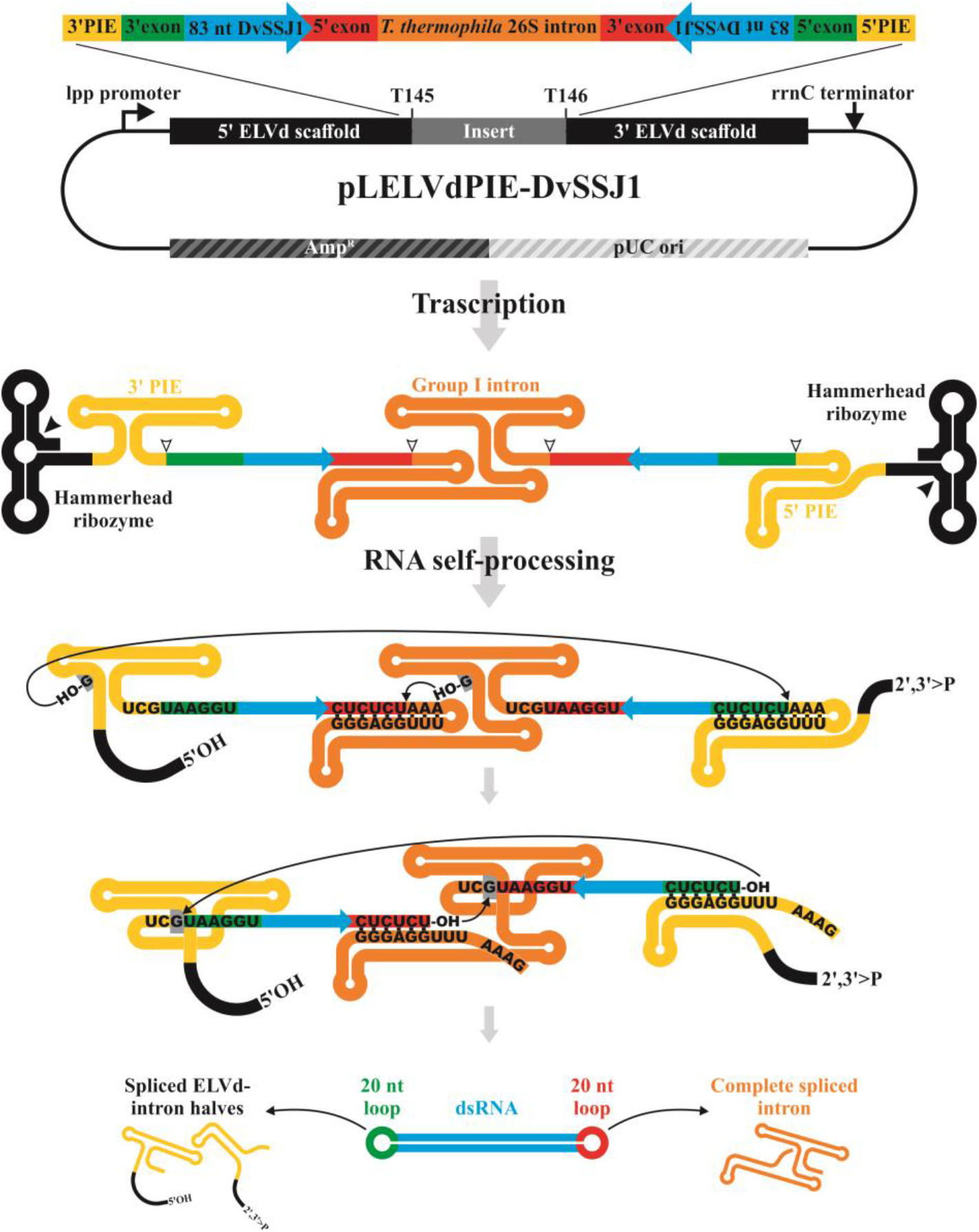
Schematic representation of the double-intron strategy to produce recombinant circular dsRNA in which the ELVd scaffold is removed. A permuted additional copy of the autocatalytic intron is added to the features of the pLELVd-DvSSJ1 flanking the inverted repeats to generate pLELVdPIE-DvSSJ1 (not at scale). Two self-splicing reactions are carried out in parallel (following the same two sequential transesterification reactions detailed in Figure 3). As a result, the complete intron is released, in addition to the two halves of the permuted intron covalently linked to the 3’ or 5’ ELVd scaffold; a circular dsRNA molecule, consisting of a 83-bp *DvSSJ1* dsRNA capped at both ends by the exon fragments is produced.

The two intron halves were amplified by PCR and inserted into the right places by the Gibson assembly reaction to build plasmid pLELVdPIE-DvSSJ1 (Supplemental Dataset S1). We co-electroporated the *E. coli* RNase III-deficient strain containing this new plasmid together with p15LtRnlSm, the plasmid to express eggplant tRNA ligase. Following plate selection of transformed clones, four independent colonies were grown in TB media for 24 h. Total bacterial RNA was extracted using phenol:chloroform and analyzed using denaturing PAGE. The controls included bacteria co-electroporated with p15LtRnlSm and either the empty ELVd plasmid (pLELVd) or pLELVd-DvSSJ1. Electrophoretic analysis of RNA preparations from bacteria transformed with pLELVdPIE-DvSSJ1 showed a new prominent band between the 100-nt and 200-nt RNA markers, which was absent in both controls (Figure 5(a), compare lanes 9 to 12, see blue arrow, with lanes 1 to 8). Surprisingly, the RNA molecule producing this band exhibited a differential migration depending on the position in the gel, creating an inverted smile pattern (Figure 5(a), lanes 9 to 12, blue arrow). This anomalous electrophoretic behavior is expected for a very compact circular dsRNA molecule, whose denaturation degree, and consequent electrophoretic mobility, changes with temperature. In this kind of electrophoresis, the temperature in the center of the gel is higher than at the sides, as is the degree of denaturation. Consequently, a compact circular dsRNA migrates at the side of the gel (less denaturing) more rapidly than it does at the center (more denaturing) (Supplemental Figure S4).

To further confirm the circularity of the recombinant dsRNA product, we subjected equivalent aliquots of RNA preparations from bacteria transformed with pLELVd and pLELVdPIE-DvSSJ1 to 2D PAGE separation. First, we separated the RNA using PAGE under non-denaturing conditions. We detected a prominent band close to the 200-bp DNA marker in the pLELVdPIE-DvSSJ1 sample; this was also present in the pLELVd control (Figure 5(b), compare lanes 1 and lane 3). However, when we split both bands, directly loaded them side-by-side on top of a second denaturing polyacrylamide gel (containing 8 M urea), and continued electrophoresis, we observed a differential band corresponding to a species with a delayed electrophoretic mobility. This arose exclusively in the half lane corresponding to ELVdPIE-DvSSJ1 (Figure 5(b), lane 4, blue arrow). These results indicate that, in bacteria transformed with pLELVdPIE-DvSSJ1, both introns (the regular and the permuted) self-splice efficiently to form a circular dsRNA product in which the ELVd scaffold has been removed, as depicted in the scheme in Figure 4.

**Figure 5.**
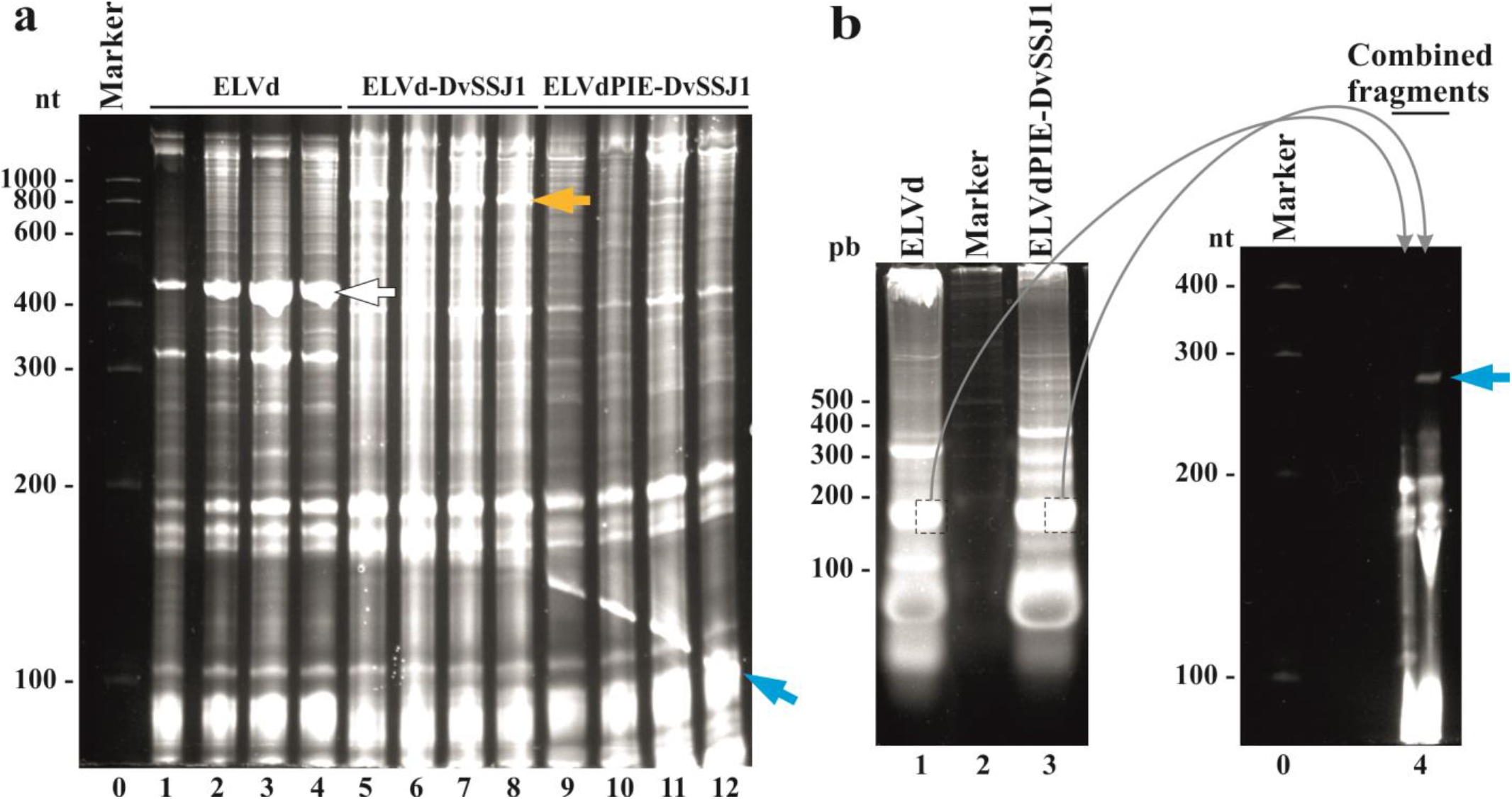
Recombinant circular dsRNA production in *E. coli* with the two-intron strategy. (a) *E. coli* HT115(DE3) was co-electroporated with p15LtRnlSm along with pLELVd (lanes 1 to 4), pLELVd-DvSSJ1 (lanes 5 to 8), or pLELVdPIE-DvSSJ1 (lanes 9 to 12). Bacterial RNAs were extracted from culture aliquots after 24 h and separated by denaturing PAGE. The gel was stained with ethidium bromide. (b) Equivalent aliquots of the total RNAs from *E. coli* co-transformed with p15LtRnlSm and either pLELVd (lane 1) or pLELVdPIE-DvSSJ1 (lane 3) were subjected to 2D PAGE separation. The first dimension (left) was conducted under non-denaturing conditions. Lane 2, DNA marker with sizes in bp on the left. The indicated bands were recovered from the first gel and placed together in a single well of a second denaturing gel (right; lane 4). The second dimension was done under denaturing conditions. (a and b) Lanes 0, RNA marker with sizes in nt on the left. Gels were stained with ethidium bromide. White, orange, and blue arrows point to ELVd, ELVd-DvSSJ1, and the circular dsRNA, respectively.

### *Circular dsRNA is also produced in* E. coli *in the absence of eggplant tRNA ligase*

Since the viroid scaffold is very effectively removed *in vivo* by the self-catalytic reactions of both introns, we examined whether the eggplant tRNA ligase added any benefit to the process. We grew cultures of bacteria co-transformed with pLELVdPIE-DvSSJ1 and either p15LtRnlSm or the corresponding empty plasmid (p15CAT; Supplemental Dataset S1). Electrophoretic analysis of the RNA preparations showed that the recombinant circular dsRNA was produced in the presence or absence of the eggplant tRNA ligase (Figure 6(a), see bllue arrow and compare lanes 1 to 3 with lanes 4 to 6).

**Figure 6.**
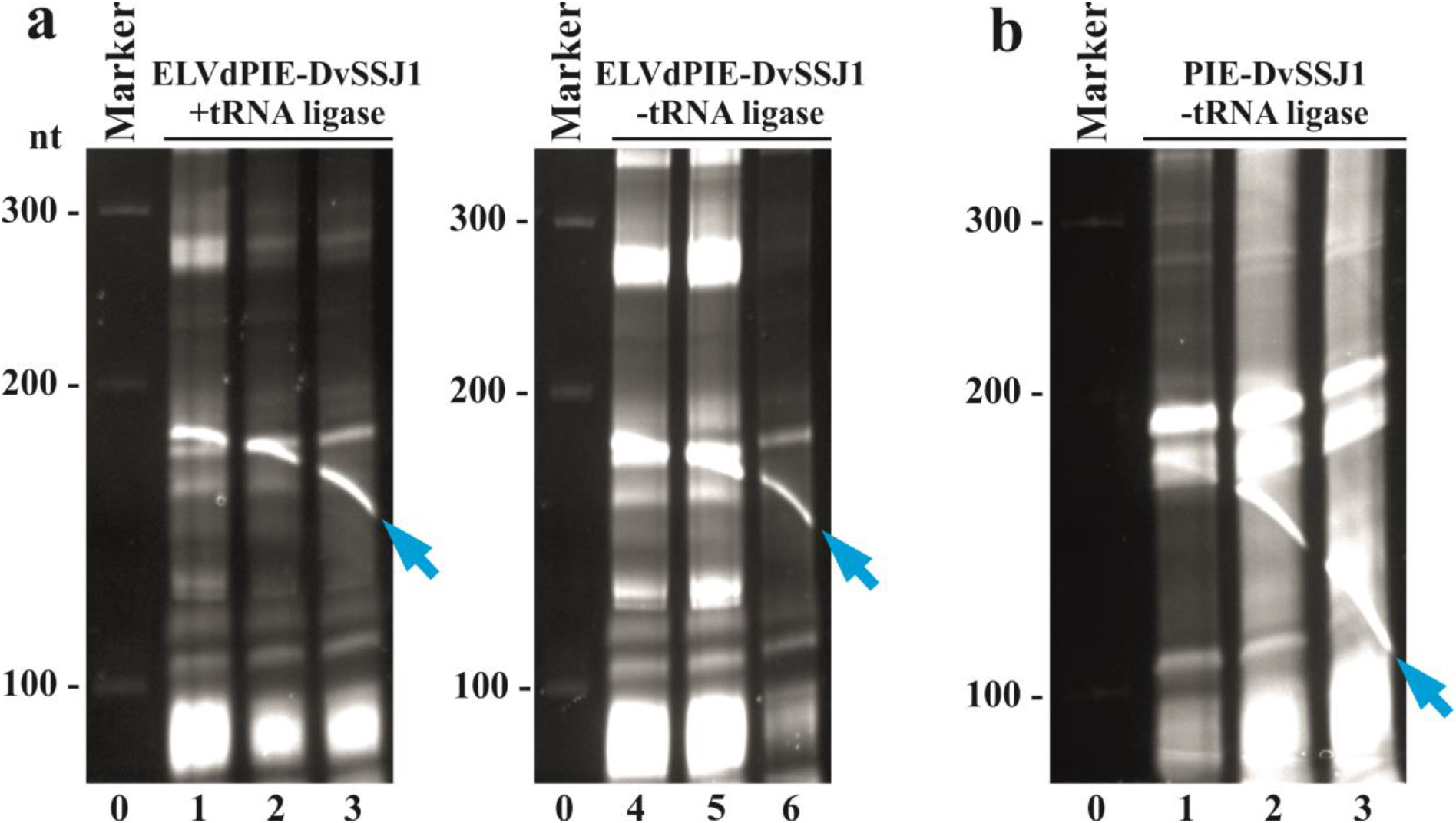
Recombinant circular dsRNA production in *E. coli* without tRNA ligase and the ELVd scaffold. RNA preparations from (a) bacteria transformed with pLELVdPIE-DvSSJ1 and p15LtRnlSm (lanes 1 to 3) or the corresponding empty plasmid p15CAT (lanes 4 to 6), and (a) bacteria transformed with pLPIE-DvSSJ1 and p15CAT (lanes 1 to 3) were separated by denaturing PAGE and the gels were stained with ethidium bromide. Lanes 0, RNA marker with sizes in nt on the left. In all cases, RNA was extracted from bacteria growing in liquid cultures after 24 h. The circular dsRNA is indicated with blue arrows.

Since this result demonstrated that the tRNA ligase was no longer required to produce the recombinant circular dsRNA, we wondered whether the ELVd scaffold itself was required. To investigate this, we constructed a new plasmid in which the two ELVd moieties were deleted (pLPIE-DvSSJ1; Supplemental Dataset S1). The RNase-III-deficient *E. coli* strain was transformed with this plasmid alone and the RNA was purified from bacteria growing in a liquid culture. Electrophoretic analysis showed that, under these new conditions with the two introns, deletion of the ELVd scaffold had only a minor effect on circular dsRNA production (Figure 6(b), blue arrow; and Supplemental Figure S5).

## Discussion

RNAi can be driven by endogenous or exogenous RNA molecules that are processed in eukaryotic cells by RNase-III-type enzymes (Dicer), resulting in small double-stranded molecules from 21 to 25 bp with two protruding nucleotides at each 3’ end. One of the strands is loaded by an Argonaute protein to form the RNA-induced silencing complex (RISC), which serves as a guide for searching RNA targets based on base complementary. Target RNAs can be processed in various ways, but the whole mechanism typically results in reduced gene expression [36]. Due to its mechanistic simplicity, RNAi has become a powerful tool for basic research and biotechnological applications. While transgenic technologies may be ideal for inducing RNAi, stringent legislation regarding genetically modified organisms makes it necessary to consider exogenously supplied RNA molecules in many instances. However, possibly due to its intrinsically low half-life, there is no easy way to produce the large amounts of recombinant RNAs that are required for many practical applications, although substantial advances in RNA expression have been made using stable RNA scaffolds in *E. coli* [8,37,38]. We have recently contributed to this effort with a system that expresses the RNA of interest in *E. coli* in a circular form that is grafted into an extremely stable scaffold consisting of a viroid RNA backbone [18]. Viroids are plant infectious agents constituted by a relatively small naked circular RNA able to survive and replicate in the hostile environment of the host cell. The circular, highly base-paired viroid structure—likely bound to the co-expressed tRNA ligase— provides the recombinant RNA with high stability and exonuclease resistance, contributing to the vast accumulation (tens of milligrams per liter of *E. coli* culture in laboratory conditions) of the RNA of interest in this system. However, this system proved to be inefficient for producing the long dsRNAs that are preferred for inducing RNAi in many applications [18].

In this work, we aimed to adapt the viroid-based system for producing recombinant RNA in *E. coli* to generate the dsRNAs that are used for RNAi-based pest control. As a target, we chose *D. virgifera*, a relevant corn pest for which RNAi-based control has been recently demonstrated by targeting the *DvSSJ1* gene [28–30]. To express a dsRNA homologous to a fragment of the *DvSSJ1* gene in the viroid-based system, we had to create a long inverted repeat in the sequence of the vector. This resulted in the longstanding and well-known problem of plasmid instability [14]. The use of asymmetric inverted repeats [9,11], or the insertion of long spacers between the repeats [10,12], has been proposed to avoid plasmid instability in bacteria. However, in other systems such as plants or *Drosophila melanogaster*, the repeats have been successfully separated with intron cDNAs from genes (of the same or related organisms) containing short fragments of both flanking exons. In this way, once transcribed, the introns are recognized and processed by the splicing machinery, eliminating them from the final product: a hairpin consisting of a long dsRNA capped by a short loop resulting from the flanking exon fragments [31,39–41]. Inspired by these studies, we separated the inverted repeats with the sequence corresponding to a self-splicing group-I intron from *T. thermophila*. Group I introns are found in genes encoding proteins, rRNA, and tRNA in algae, fungi, lichens, some lower eukaryotes, and especially in bacteria. These introns are ribozymes, capable of catalyzing their own splicing from primary transcripts without the involvement of any protein or additional factor, other than Mg^2+^ and guanosine [42]. In addition to the full-length cDNA of the *T. thermophila* 26S rRNA intron (433 nt), we added each 10-nt flanking exon to ensure optimal recognition of intron-exon boundaries. As expected, the separation of the inverted repeats with the intron plus flanking exons was key to achieving plasmid stability in *E. coli* (Supplemental Figure S1). The expression of the corresponding precursor transcript in an *E. coli* RNase III-deficient strain, along with that of the eggplant tRNA ligase, led to the efficient accumulation of chimeric molecules of circular RNA consisting of the ELVd scaffold containing the 83-nt *DvSSJ1* hairpin (Figure 1). Interestingly, the 433-nt processed intron also accumulated in *E. coli*, while unprocessed intermediates were not detected, indicating very efficient *T. thermophila* group-I intron self-cleavage in bacteria (Figure 2) [43]. The amount of the chimeric RNA containing the dsRNA of interest increased with time up to 24 h in the *E. coli* culture, in contrast to the usual 12-h accumulation peak of single-stranded RNA aptamers, as in Spinach [18]. The amount decreased after that point, with the band of circular RNA at 40 h being almost negligible (Figure 2). In addition, the 50-fold upscaling of production (maintaining the expression conditions) does not seem to affect the large-scale accumulation of the dsRNA of interest. Thus, we suggest that this approach is an efficient way to produce large amounts of recombinant dsRNAs, due to the fact that both the scaffolding and the hairpin loop are constructed to avoid degradation by the bacterial RNases.

The produced dsRNA maintains a potent anti-WCR activity, as we observed in the WCR larvae feeding bioassay (Table 1). As previously shown, the inserted RNA is fully functional, probably due to the position in which the recombinant RNA is inserted—at the end of the right upper arm, thus allowing both scaffold and recombinant RNA to form two independent domains [18]. Furthermore, it should be noted that the silencing obtained with the ELVd-dsRNA chimera is slightly better than that obtained using the same dsRNA molecules produced by *in vitro* transcription (Table 1). We speculate that the presence of the protective scaffold may help increase the half-life of dsRNA during its production and application, both in the field and in the intestines and cells of insects, thereby making it available for processing by the RNAi machinery in greater quantities than the same dsRNA produced *in vitro*.

Various releasing methods have been described to remove the protective scaffold after expression of recombinant RNA. *In vitro* strategies such as the use of RNases [10,44] or sequence-specific DNAzymes [37], although they remove single-stranded RNA, would add an additional layer to the production system. Furthermore, the resulting RNA would not be circular. Another feasible approach is to flank the recombinant RNA ends with hammerhead ribozymes, which leads to *in vivo* efficient cleavage of the RNA of interest and its release as a linear [45] or even a circular molecule [46]. In this work, we explore the use of an additional copy of the same group-I autocatalytic intron, although in a permuted fashion in which the 3’ half of the intron is placed upstream of the 5’ half (Figure 4). As reported, both intron halves undergo the two transesterification reactions [35], even inserting between both halves very long sequences [47] such as, in this case, the long inverted repeats plus the full-length separating intron. The resulting product is a circular molecule composed of the dsRNA locked at both sides by loops derived from the exon fragments (Figure 4). This strategy allows us to remove the viroid scaffolding without losing, in the final molecule, two key characteristics in recombinant RNA accumulation—circularity and compaction—that make the molecules highly resistant to degradation, as seen from the fact that they accumulate in large quantities in the bacteria (Figure 5). Interestingly, although there are two complete intron sequences, they are both removed correctly without mutual interference. Due to the efficient self-splicing of both introns, the circular dsRNA can be efficiently obtained in *E. coli*, even if neither the eggplant tRNA ligase nor the viroid scaffold are present (Figure 6).

Purified dsRNA or even inactivated complete bacterial extracts have been used to induce RNAi effectively [15,16]. For many applications, it is preferable, however, to completely isolate the dsRNA from other bacterial RNAs. Thus, both recombinant dsRNAs produced here could be purified by affinity chromatography, using recombinant-specific dsRNA-binding proteins or antibodies [48]. The circularity of the produced molecules and the absence of such structures in *E. coli* may also be exploited, as the circular molecules can be purified to homogeneity by 2D-PAGE, both under denaturing conditions (with reduction of the ionic strength in the second separation) and subsequent elution from the gel.

In conclusion, we have developed a strategy to adapt our viroid-based expression system to overproduce in *E. coli* RNA hairpins with extended double-stranded regions able to induce RNAi in insects, as we demonstrate in the case of WCR. Further, we used a second self-splicing group-I intron (permuted) to produce very compact circular dsRNAs in which the viroid scaffold is completely removed *in vivo*. Both strategies are based on the activity of autocatalytic introns. We assert that they are fast, high-output, cost-effective, and scalable alternatives for industrially producing the large amounts of large dsRNAs that are required in pest control approaches. Because of their robustness and flexibility, these strategies could be used in any other RNAi biotechnological applications.

## Materials and methods

### Plasmid construction

To build pLELVd-DvSSJ1, we first produced an 83-bp cDNA molecule homologous to a fragment of *DvSSJ1* (from positions 50 through 132 of GenBank KU562965.1) via the polymerase chain reaction (PCR), using the Phusion high-fidelity DNA polymerase (Thermo Scientific) and primers D2623 and D2624. All primers used in this work are in Supplemental Table S1. Next, we amplified three cDNAs corresponding to this *DvSSJ1* fragment in two opposite orientations, using primers D2625-D2626 and D2629-D2630, as well as the *T. thermophila* 26S rRNA intron, which includes 10 nt of the flanking exons (from positions 43 through 475 of V01416.1) with primers D2627 and D2628. Finally, we assembled [49] these three cDNAs into pLELVd-BZB (Supplemental Dataset S1), digested with *Bpi*I (Thermo Scientific). The sequence of the resulting plasmid, pLELVd-DvSSJ1 (Supplemental Dataset S1), was experimentally confirmed (3130xl Genetic Analyzer, Life Technologies). From this plasmid, we built pLELVdPIE-DvSSJ1 by adding, through two consecutive Gibson assembly reactions, two halves of the autocatalytic intron: 3’ (opening the plasmid with the D2936 and D2937 primers) and 5’ (opening the plasmid with the primers D2940 and D2941). We also sequentially removed the 5’ and 3’ viroid moieties from this latter plasmid via PCR with the phosphorylated primers (T4 polynucleotide kinase, Thermo Scientific) D3606 and D3285, and D3607 and D3608, followed by self-ligation of the products (T4 DNA ligase, Thermo Scientific). Finally, we obtained plasmid pLPIE-DvSSJ1 (Supplemental Dataset S1).

### Escherichia coli *culture*

The strain HT115(DE3) [15] of *E. coli* was co-electroporated (Eporator, Eppendorf) with p15LtRnlSm along with pLPP, pLELVd, pLELVd-DvSSJ1, pLELVdPIE-DvSSJ1, or pLPIE-DvSSJ1. In some experiments, bacteria were electroporated only with pLELVdPIE-DvSSJ1 or pLPIE-DvSSJ1. Transformed clones were selected at 37ºC in plates of Luria-Bertani (LB) solid medium (10 g/l tryptone, 5 g/l yeast extract, 10 g/l NaCl, and 1.5% agar) that included the appropriate antibiotics (50 μg/ml ampicillin and 34 μg/ml chloramphenicol, when needed). Liquid cultures of *E. coli* were grown in TB medium (12 g/l tryptone, 24 g/l yeast extract, 0.4% glycerol, 0.17 M KH_2_PO_4_, and 0.72 M K_2_HPO_4_), which also contained the appropriate antibiotics (as above), at 37ºC with vigorous shaking (225 revolutions per min; rpm).

### RNA extraction

At the desired time, 2-ml aliquots of the liquid cultures were harvested; the cells were sedimented by centrifugation at 13,000 rpm for 2 min. Cells were resuspended in 50 μl of TE buffer (10 mM Tris-HCl, pH 8.0 and 1 mM EDTA). One volume (50 μl) of a 1:1 (v/v) mix of phenol (saturated with water and equilibrated at pH 8.0 with Tris-HCl, pH 8.0) and chloroform was added; the cells were broken by vigorous vortexing. The mix was centrifuged for 5 min at 13,000 rpm; the aqueous phase, containing total bacterial RNA, was recovered. For large-scale preparation of ELVd and ELVd-DvSSJ1 RNA, *E. coli* were grown in a 2-l Erlenmeyer flask with 250 ml of TB medium at 37ºC and 225 rpm for 24 h. Cells were sedimented by centrifugation for 15 min at 8000 rpm, resuspended in H_2_O, and sedimented again under the same conditions. Cells were resuspended in 10 ml buffer 50 mM Tris-HCl, pH 6.5, 0.15 M NaCl, and 0.2 mM EDTA. One volume of a 1:1 mix of phenol and chloroform was added; the mix was intensively vortexed. The phases were separated by centrifugation; the aqueous phase, once recovered, was re-extracted with one volume of chloroform. Finally, RNAs were precipitated from the aqueous phase adding sodium acetate pH 5.5 to 0.3 M and 2.5 volumes of ethanol. RNAs were resuspended in H_2_O and re-precipitated with one volume of isopropanol.

### RNA electrophoresis

Aliquots of the RNA preparations (20 μl; corresponding to 0.8 ml of the original *E. coli* culture) were mixed with one volume of loading buffer (98% formamide, 10 mM Tris-HCl, pH 8.0, 1 mM EDTA, 0.0025% bromophenol blue, and 0.0025% xylene cyanol), denatured (1.5 min at 95ºC followed by snap cooling on ice), and separated by denaturing PAGE. The Riboruler low range RNA ladder (Thermo Scientific) was used as a standard. Gels were run for 2 h at 200 V in 140 × 130 × 2 mm, 5% polyacrylamide gels (37.5:1 acrylamyde:N,N’-methylenebisacrylamide) in TBE buffer (89 mM Tris, 89 mM boric acid, 2 mM EDTA) that included 8 M urea. The electrophoresis buffer was TBE without urea. The gels were stained by shaking for 15 min in 200 ml of 1 μg/ml ethidium bromide. After being washed three times with water, the gels were photographed under UV light (UVIdoc-HD2/20MX, UVITEC). In one experiment, an RNA preparation was separated by denaturing electrophoresis in a second dimension at a lower (0.25× TBE) ionic strength. After the first dimension, an entire lane from a 5% polyacrylamide, 8 M urea, TBE gel was cut and laid transversely on top of a 5% polyacrylamide gel of the same dimensions in 0.25× TBE buffer containing 8 M urea; it was run for 2.5 h at 25 mA. We set an upper limit of 25 mA. In another experiment, the RNA was separated first in a non-denaturing 5% PAGE gel in TAE buffer (40 mM Tris, 20 mM sodium acetate, 1 mM EDTA, pH 7.2) without urea. The gel was run for 1.5 h at 75 mA. The bands of interest were cut after the electrophoresis separation and placed on top of a 5% PAGE gel in 0.25× TBE buffer containing 8 M urea; they were run as previously explained.

### Northern blot hybridization analysis of RNA

After electrophoretic separation, RNAs were electroblotted to positively charged nylon membranes (Nytran SPC, Whatman) and cross-linked by irradiation via 1.2 J/cm^2^ UV light (254 nm, Vilber Lourmat). Hybridization was performed overnight at 70ºC in 50% formamide, 0.1% Ficoll, 0.1% polyvinylpyrrolidone, 100 ng/ml salmon sperm DNA, 1% sodium dodecyl sulfate (SDS), 0.75 M NaCl, 75 mM sodium citrate, pH 7.0, with approximately 1 million counts per minute of ^32^P-labelled RNA probe. Hybridized membranes were washed three times for 10 min with 2× SSC, 0.1% SDS at room temperature; they were washed again for 15 min at 55ºC with 0.1× SSC, 0.1% SDS. SSC buffer is 150 mM NaCl, 15 mM sodium citrate, pH 7.0. The results were registered by autoradiography using X-ray films (Fujifilm). Radioactive RNA probes complementary to ELVd, the *DvSSJ1* fragment, and the *T. thermophila* intron were obtained by *in vitro* transcription of the corresponding linearized plasmids with 20 U of T3 bacteriophage RNA polymerase (Roche) in 20-μl reactions containing 40 mM Tris-HCl, pH 8.0, 6 mM MgCl_2_, 20 mM DTT, 2 mM spermidine, 0.5 mM each of ATP, CTP, and GTP, and 50 μCi of [α-^32^P]UTP (800 Ci/mmol), 20 U RNase inhibitor (RiboLock, Thermo Scientific), and 0.1 U yeast inorganic pyrophosphatase (Thermo Scientific). The reactions were incubated for 1 h at 37 ºC. After transcription, the DNA template was digested with 20 U DNase I (Thermo Scientific) for 10 min at 37°C; the probe was purified by chromatography using a Sephadex G-50 column (mini Quick Spin DNA Columns, Roche).

### Double-stranded RNA production by in vitro transcription

The target-specific primers containing T7 RNA polymerase sites at the 5’ end of each primer were used to generate the PCR product; this served as the template for dsRNA synthesis by *in vitro* transcription using a MEGAscript kit (Life Technologies). The dsRNAs were purified using the Megaclear kit (Life Technologies) and examined by 12-well E-gel electrophoresis (Life Technologies) to ensure dsRNA integrity. They were quantified using Phoretix 1D (Cleaver Scientific) or a NanoDrop 8000 Spectrophotometer (Thermo Scientific).

### WCR bioassays

We prepared the WCR diet according to the manufacturer’s guideline for a *D. virgifera* diet (Frontier, Newark, DE), with modifications [50]. The dsRNA samples (5 μl) were incorporated into 25 μl of WCR diet in a 96-well microtiter plate and shaken on an orbital shaker for 1 min until the diet solidified. For each RNA sample, nine doses (35, 17.5, 8.8, 4.4, 2.2, 1.09, 0.55, 0.27, and 0.14 ng/μl) were evaluated, for a total of 32 observations per dose or water control. We transferred two one-day-old larvae into each well. The plates were incubated at 27ºC and 65% relative humidity. Seven days after exposure, the larvae were scored for growth inhibition (severely stunted larvae with >60% reduction in size) and mortality. We analyzed the data using PROC Probit analysis in SAS [51] to determine LC_50_. The total numbers of dead and severely stunted larvae were used to analyze the IC_50_.

## Disclosure statement

No potential conflict of interest was reported by the authors.

## Funding

This work was supported by the Ministerio de Ciencia e Innovación (Spain) through the Agencia Estatal de Investigación (grants BIO2017-83184-R and BIO2017-91865-EXP; co-financed by the European Region Development Fund). B.O. is the recipient of a predoctoral contract from Universitat Politècnica de València (PAID-01-17).

## SUPPLEMENTAL DATA

**Supplemental Dataset S1.**
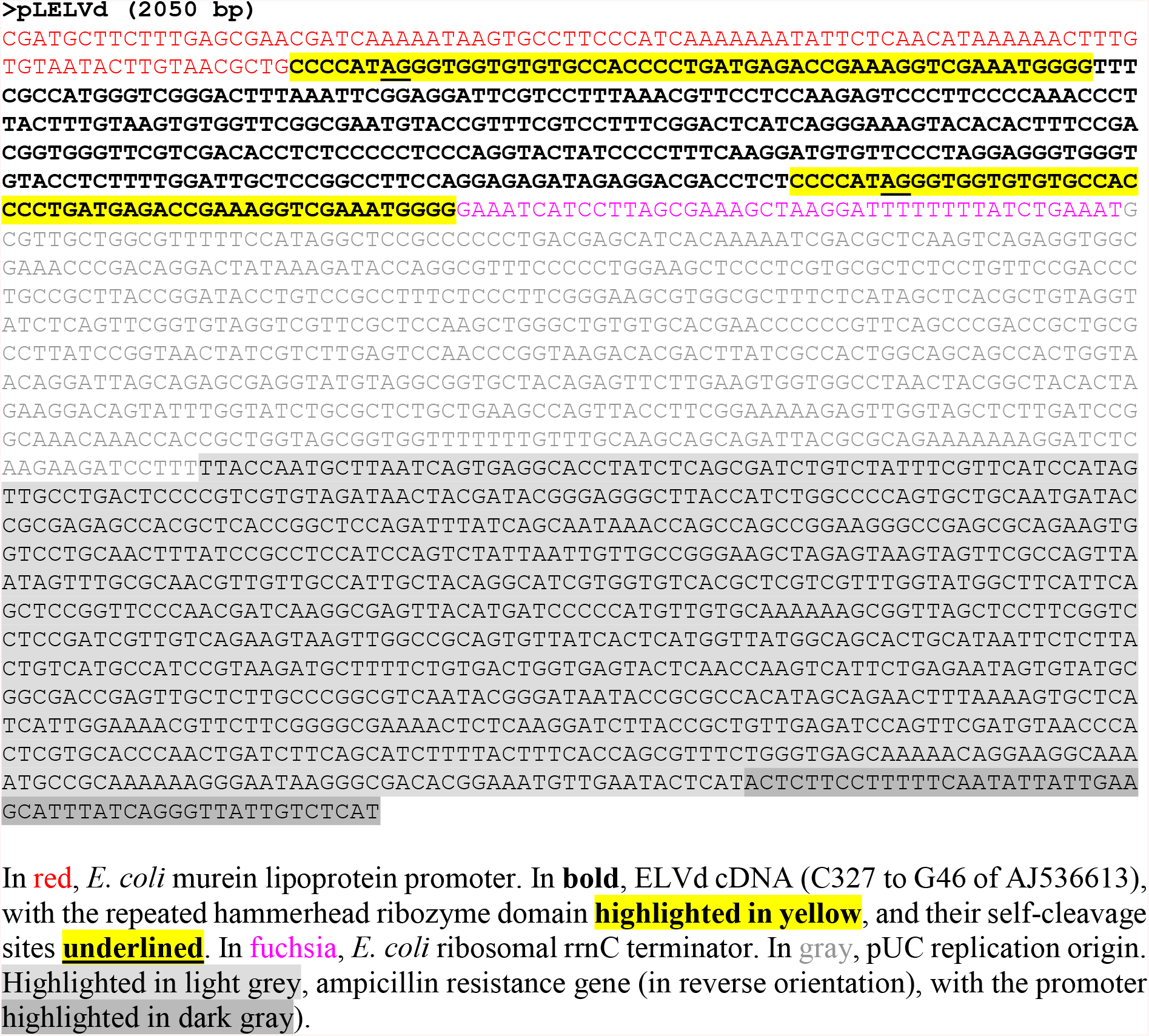
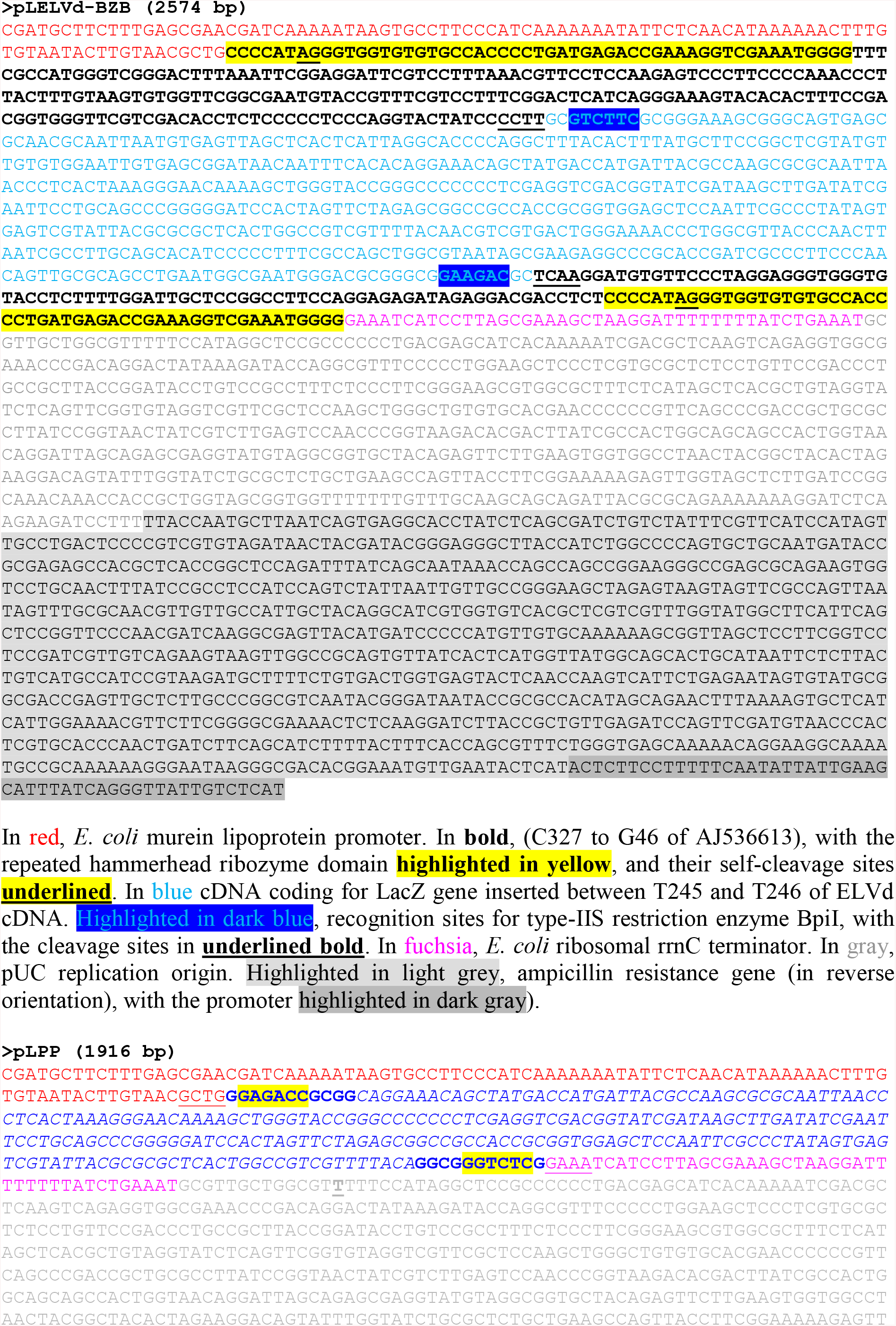
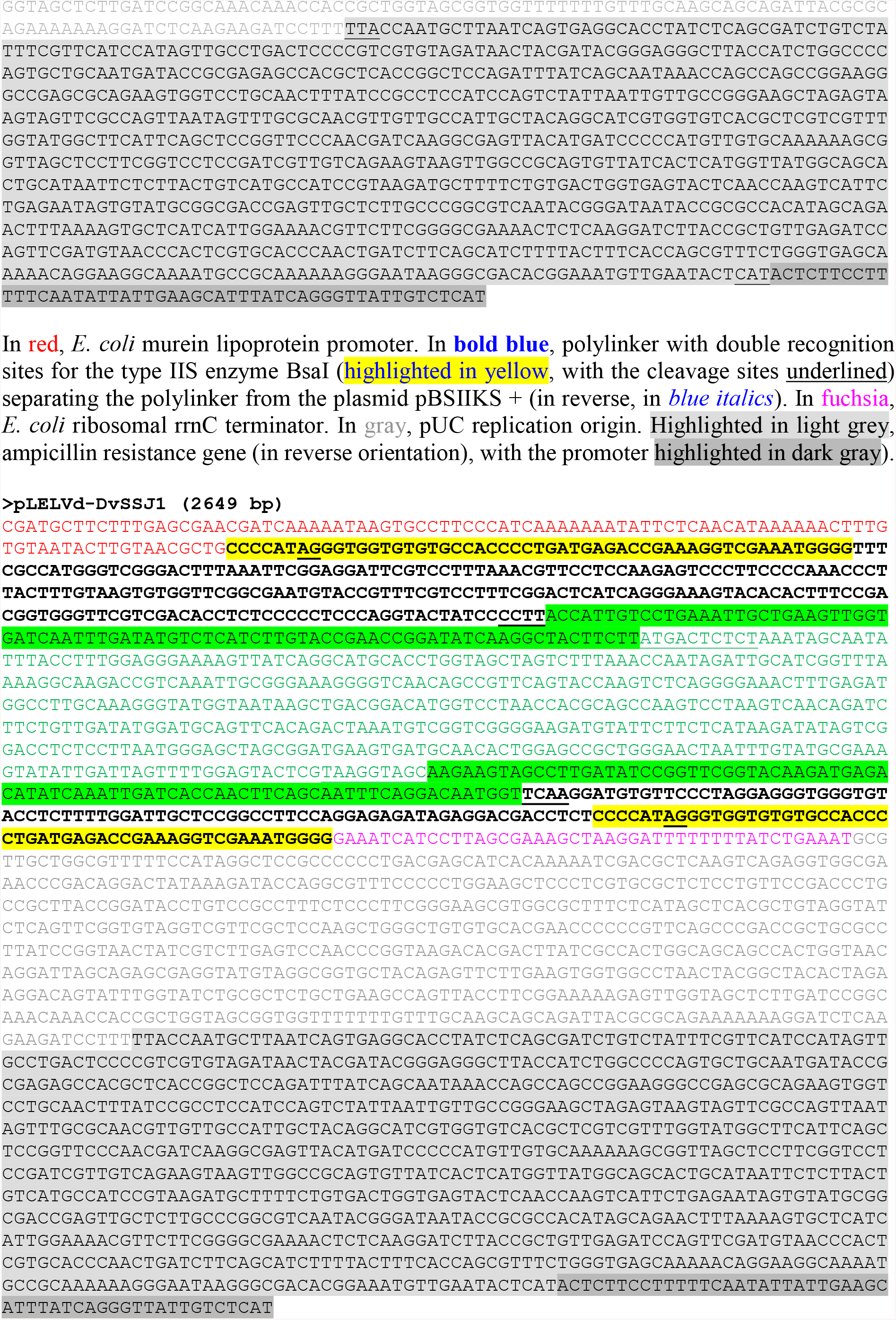
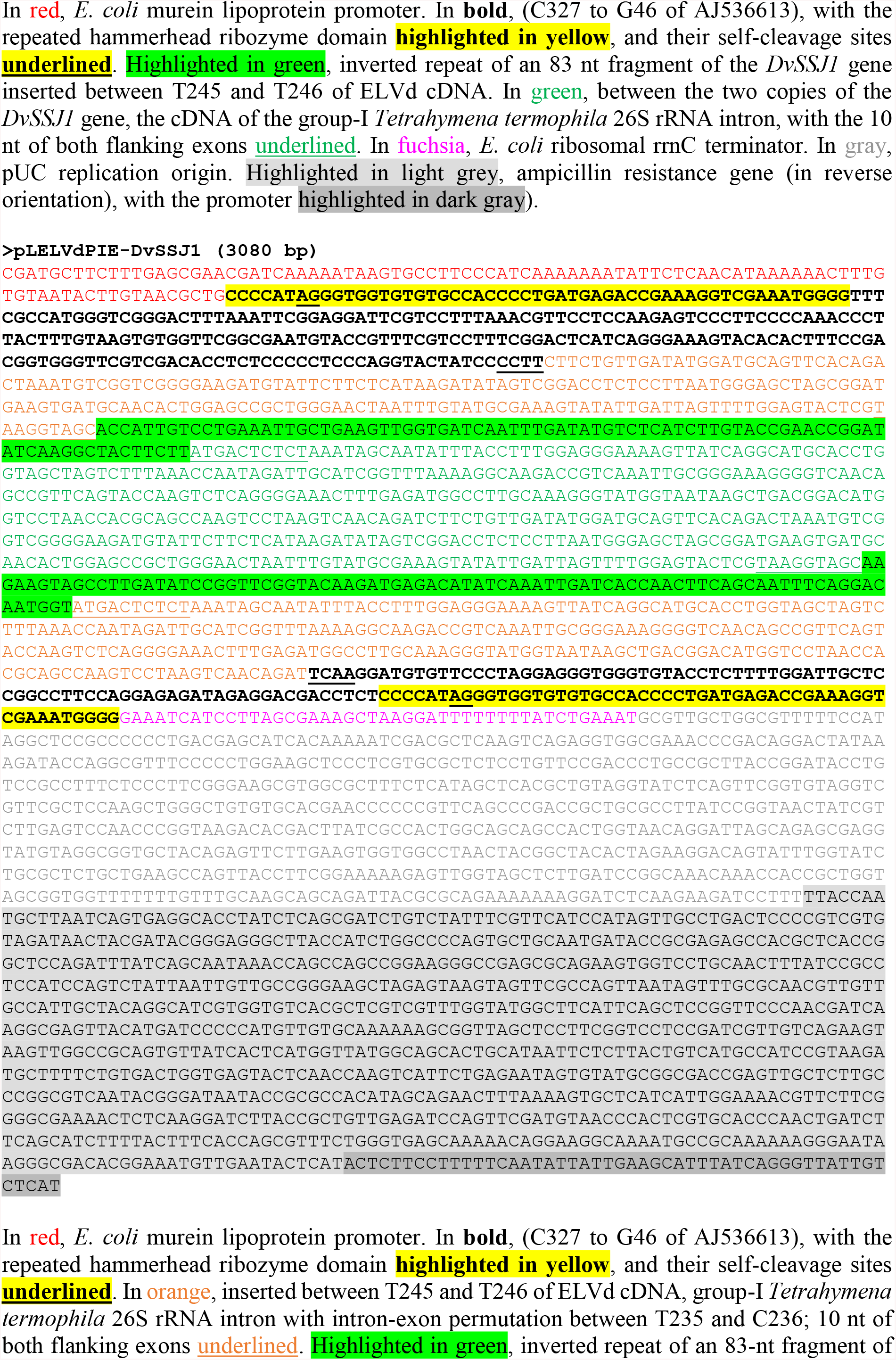
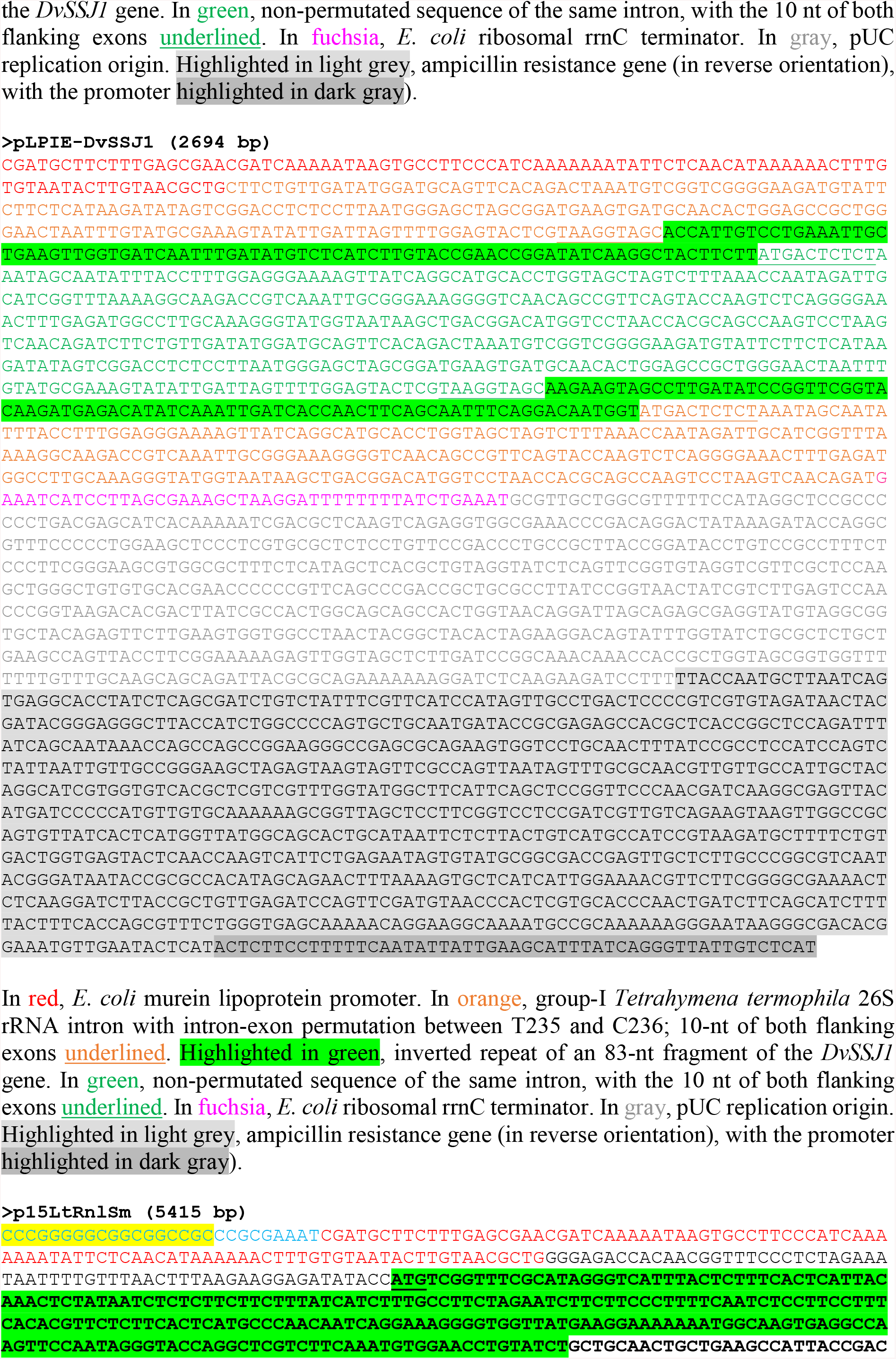
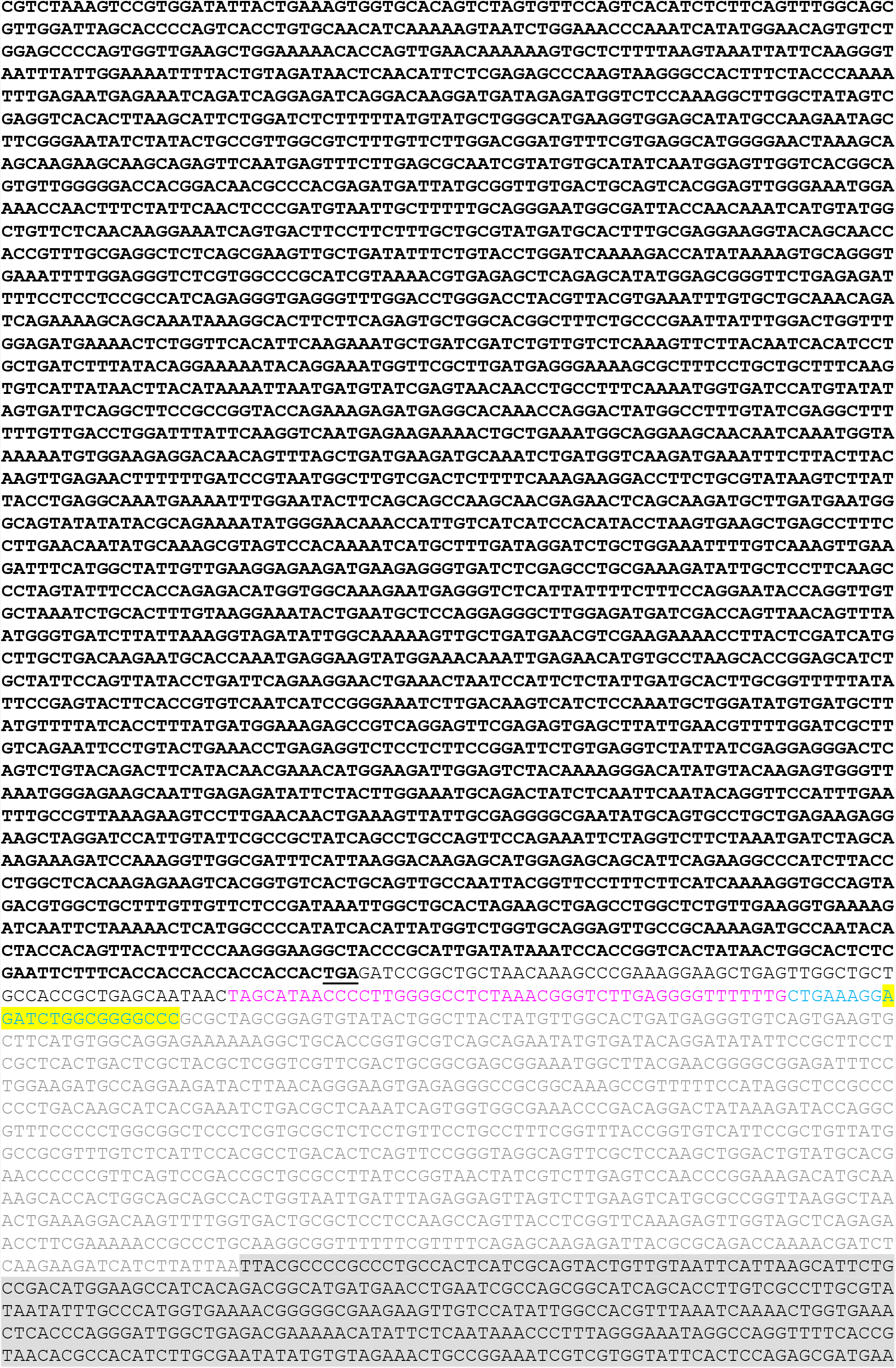
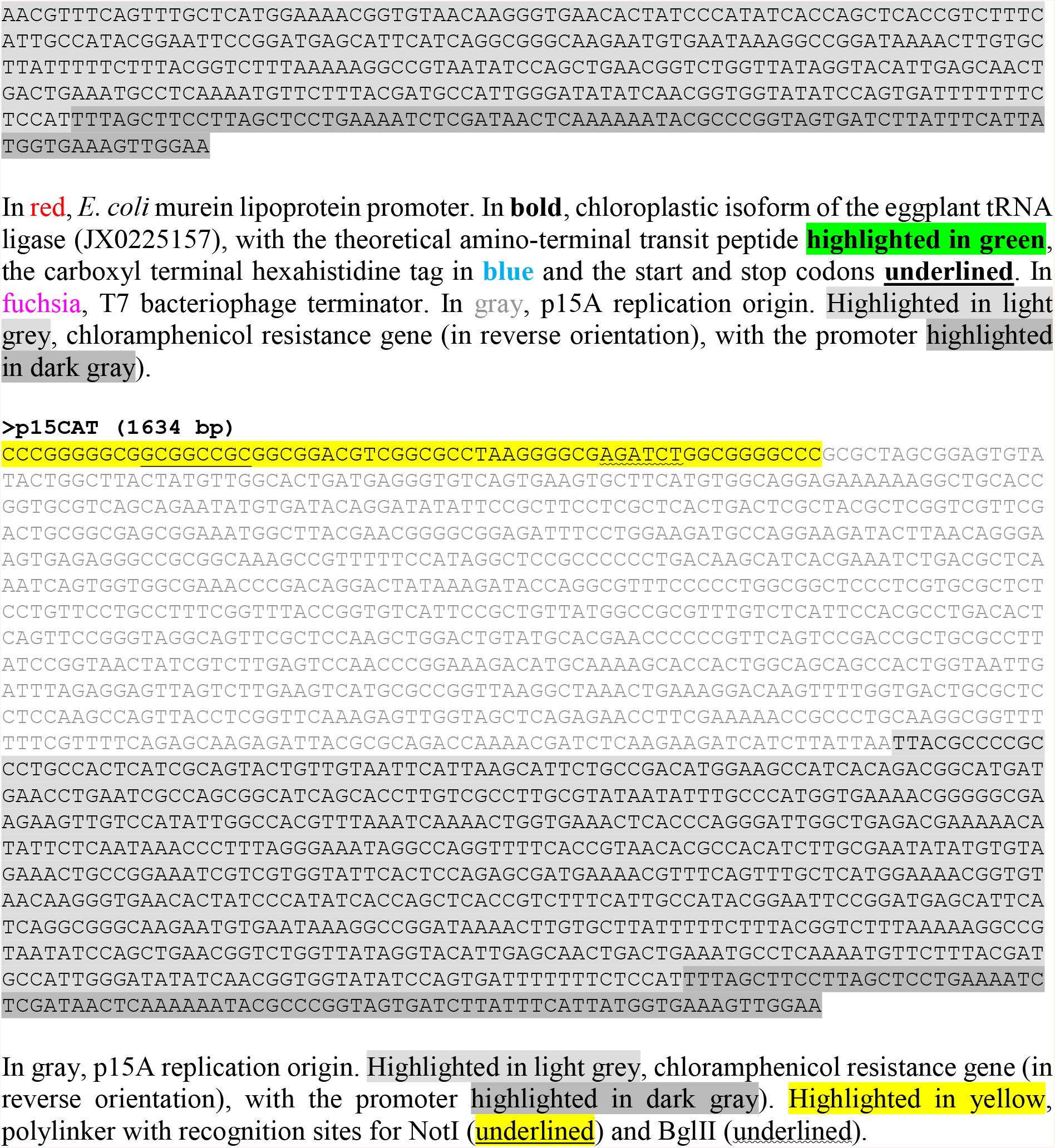
Nucleotide sequences and elements of plasmids pLELVd, pLELVd-BZB, pLPP, pLELVd-DvSSJ1, pLELVdPIE-DvSSJ1, pLPIE-DvSSJ1, p15LtRnlSm, and p15CAT.

**Supplemental Figure S1.**
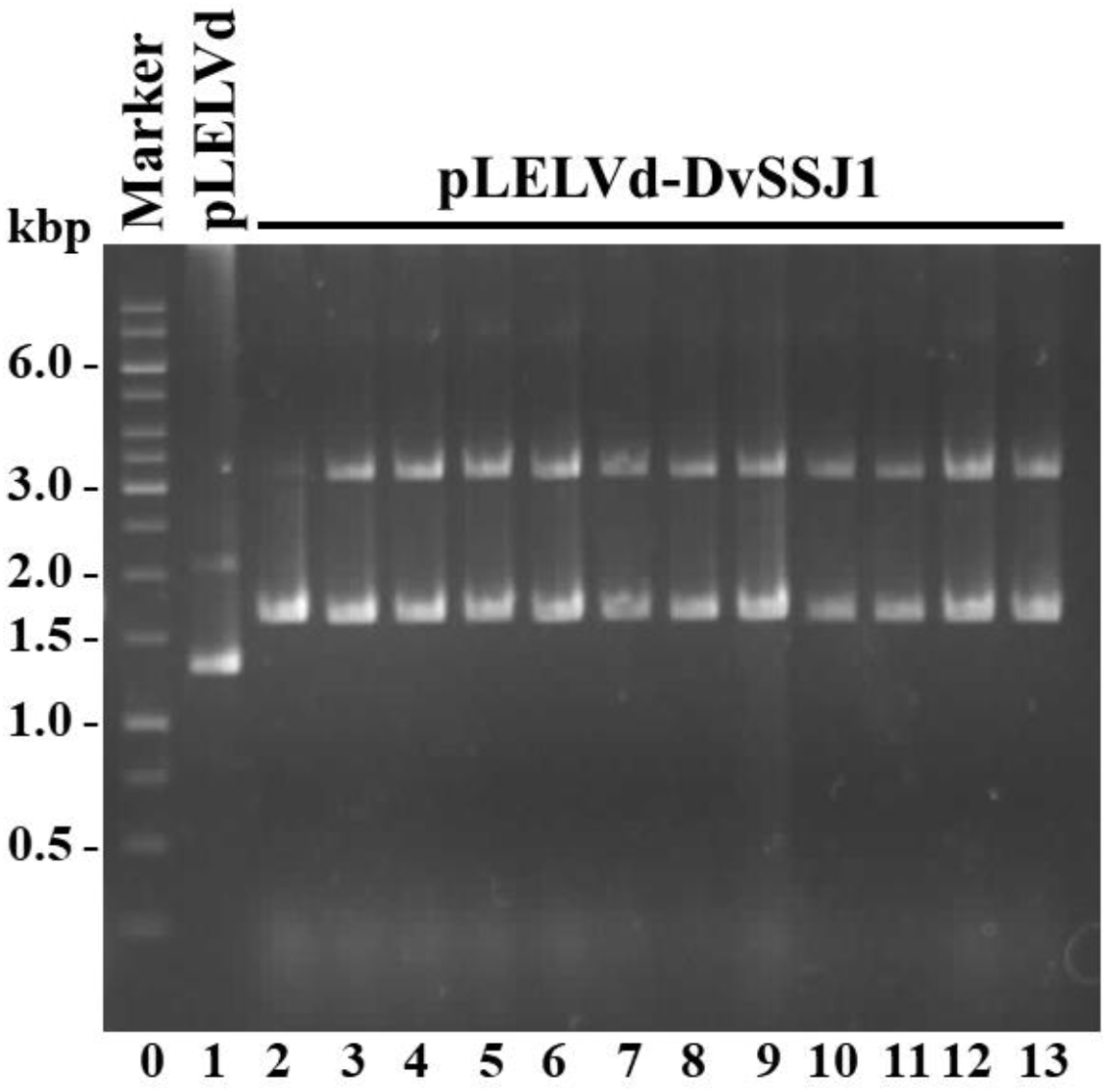
Construction of expression plasmids to produce *DvSSJ1*-derived dsRNA in *E. coli*. Plasmids purified from independent *E. coli* clones were separated by electrophoresis through an agarose gel, which was stained with ethidium bromide. Lane 0, DNA marker ladder with some of the sizes in bp on the left; lane 1, control plasmid pLELVd expressing an empty ELVd; lanes 2 to 13, plasmids pLELVd-DvSSJ1 to express the *DvSSJ1*-derived dsRNA on an ELVd scaffold obtained from 12 independent *E. coli* colonies.

**Supplemental Figure S2.**
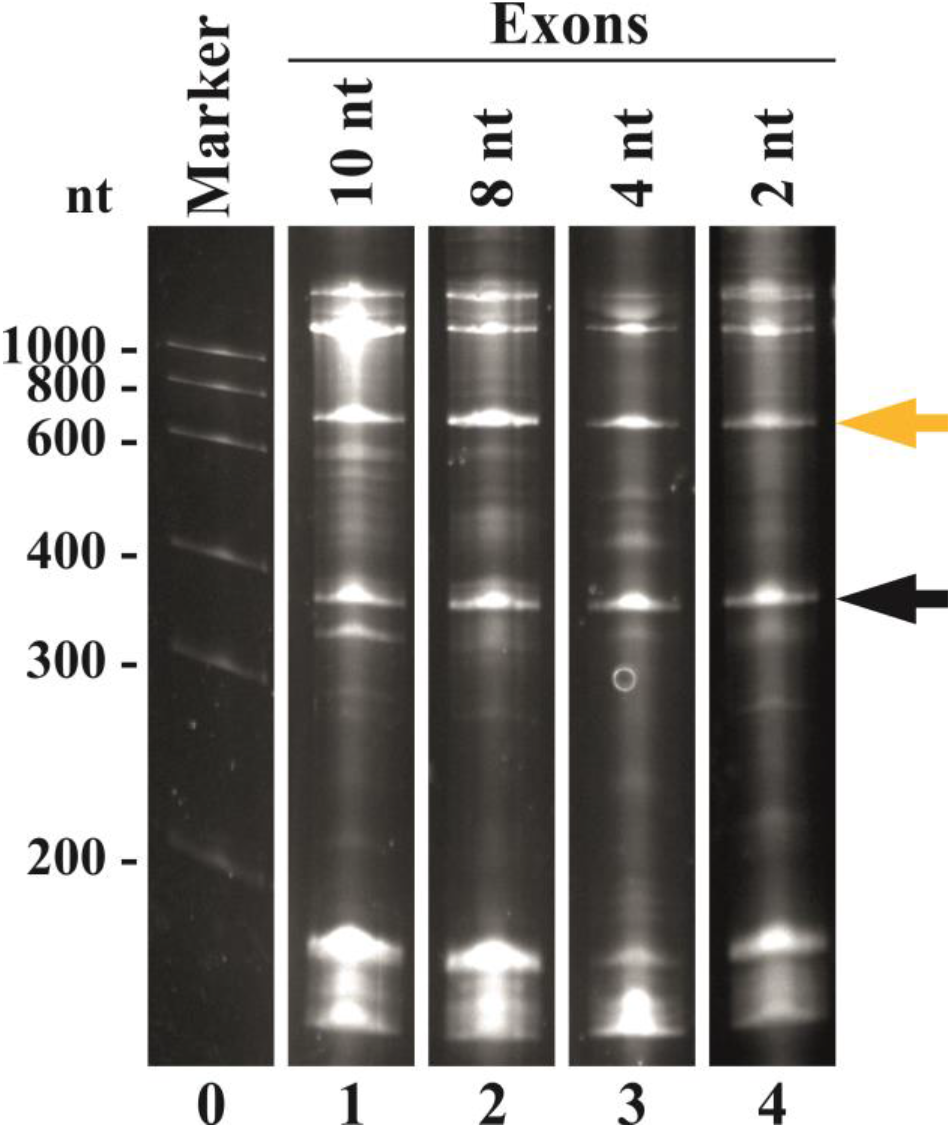
Effect of exon size in *T. thermophila* intron processing in the ELVd-based system to produce dsRNA in *E. coli*. Aliquots of RNA preparations from *E. coli* clones cotransformed with p15LtRnlSm and a series of pLELVd derivatives to produce a 100-bp dsRNA, in which the exons that flank the *T. thermophila* intron are increasingly shorter, as indicated, were separated by denaturing PAGE. The gel was stained with ethidium bromide. Lane 0, RNA marker with sizes (in nt) on the left; lanes 1 to 4, RNAs from constructs with 10, 8, 4 and 2-nt exons, respectively. Orange and black arrows point the positions of the recombinant ELVd-dsRNA and the spliced introns, respectively.

**Supplemental Figure S3.**
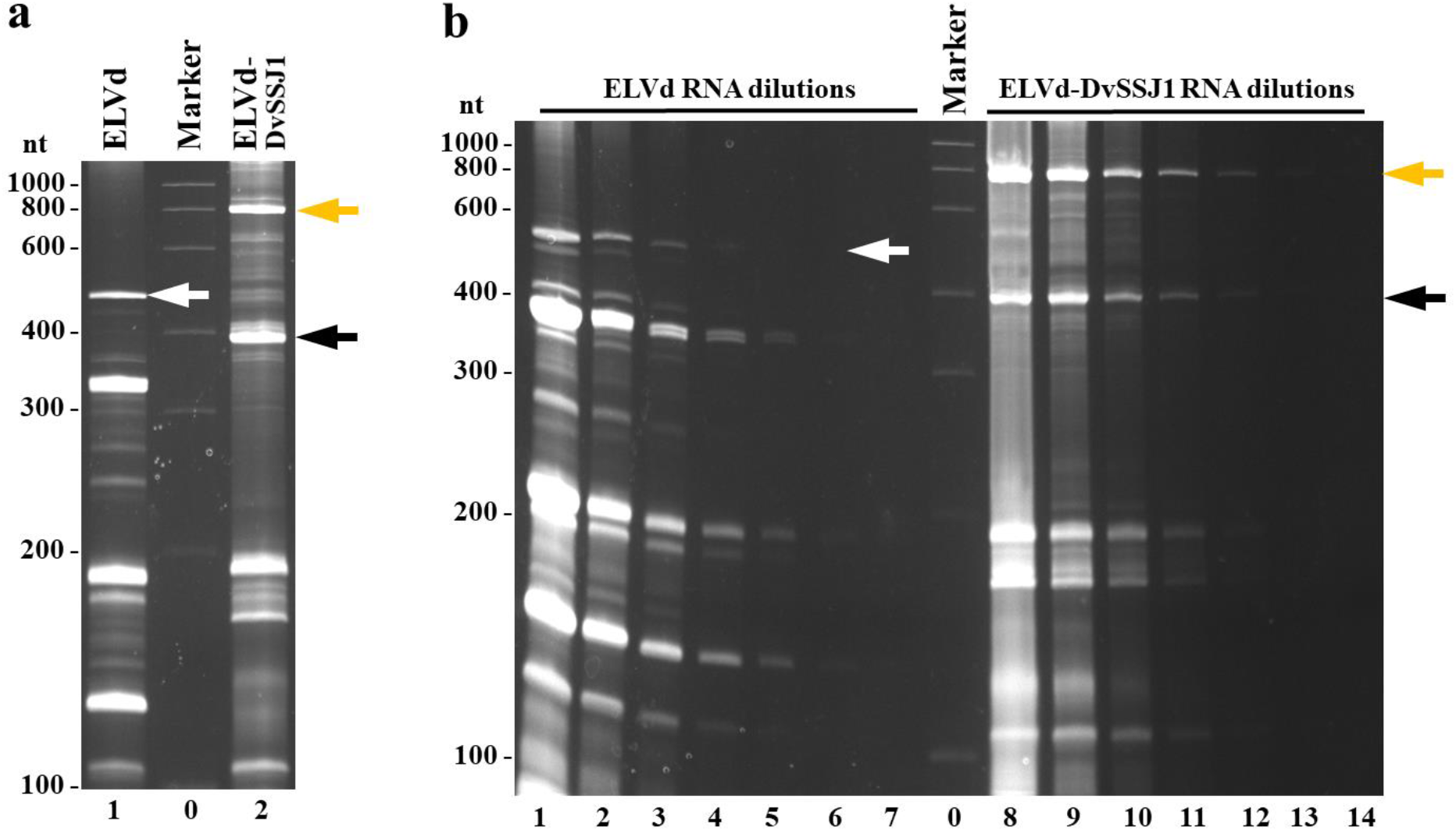
Large scale RNA preparations produced in *E. coli* by means of the viroid-based system and used in the WCR bioassay. RNAs were separated by denaturing PAGE and the gels stained with ethidium bromide. (a and b) Lane 0, RNA marker ladder with sizes in nt on the left. (a) Lanes 1 and 2, large-scale RNA preparations from *E. coli* transformed with p15LtRnlSm and pLELVd or pLELVd-DvSSJ1, respectively. (b) Dilution analysis of the ELVd (lanes 1 to 7) and the ELVd-DvSSJ1 (lanes 8 to 14) RNA preparations. White, orange and black arrows point to ELVd, ELVd-DvSSJ1 and spliced-intron RNAs, respectively.

**Supplemental Figure S4.**
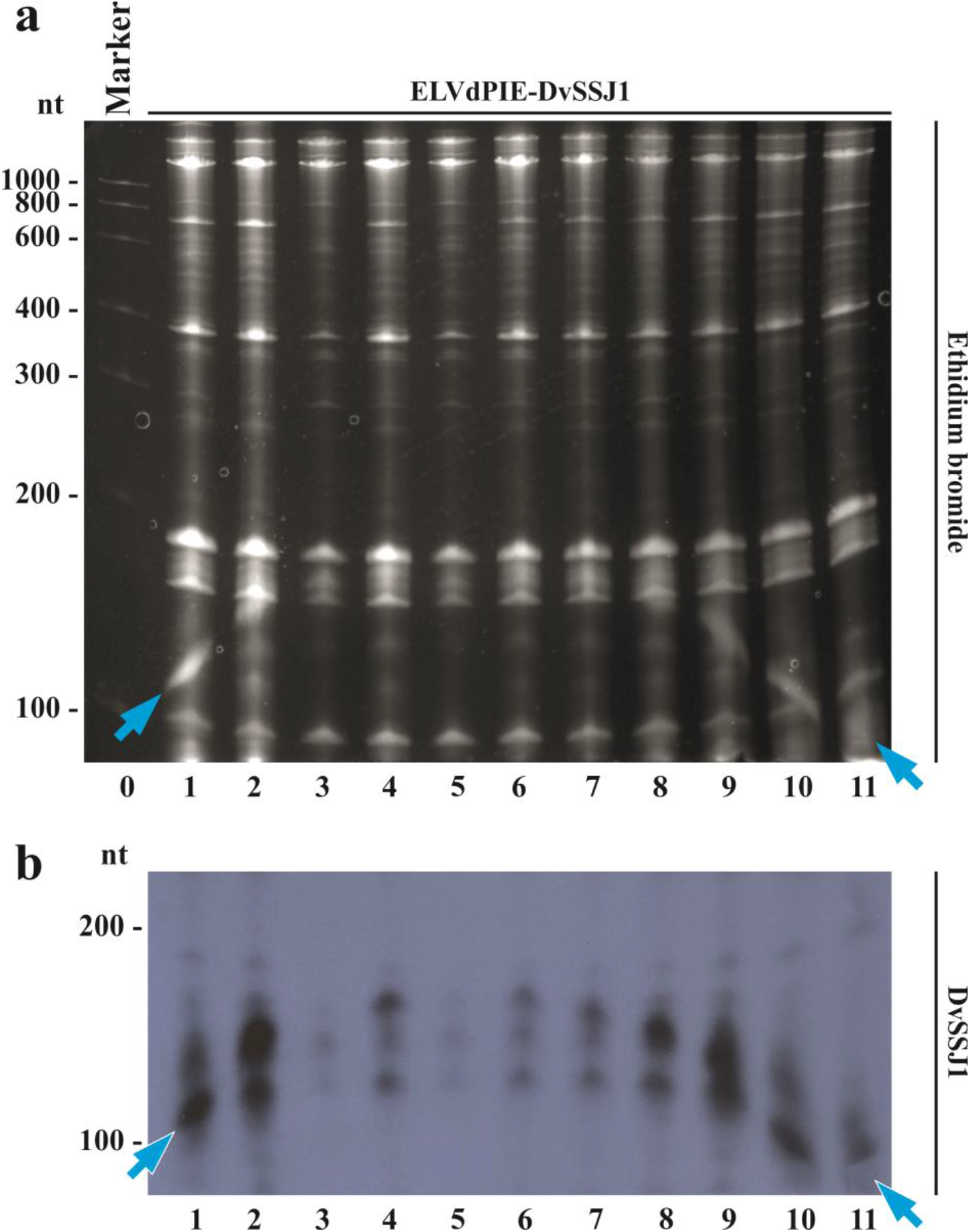
Analysis of the recombinant circular dsRNA. RNApreparations from different *E. coli* clones (lanes 1 to 11) co-transformed with p15LtRnlSm and pLELVdPIE-DvSSJ1 were separated by denaturing PAGE. The gel was (a) stained with ethidium bromide and (b) the RNA transferred to a membrane and hybridized with a ^32^P-labelled probe to detect DvSSJ1 RNA. Lane 0, RNA marker with sizes in nt on the left. Blue arrows point to the recombinant circular dsRNA that exhibits an inverted smile migration across the gel.

**Supplemental Figure S5.**
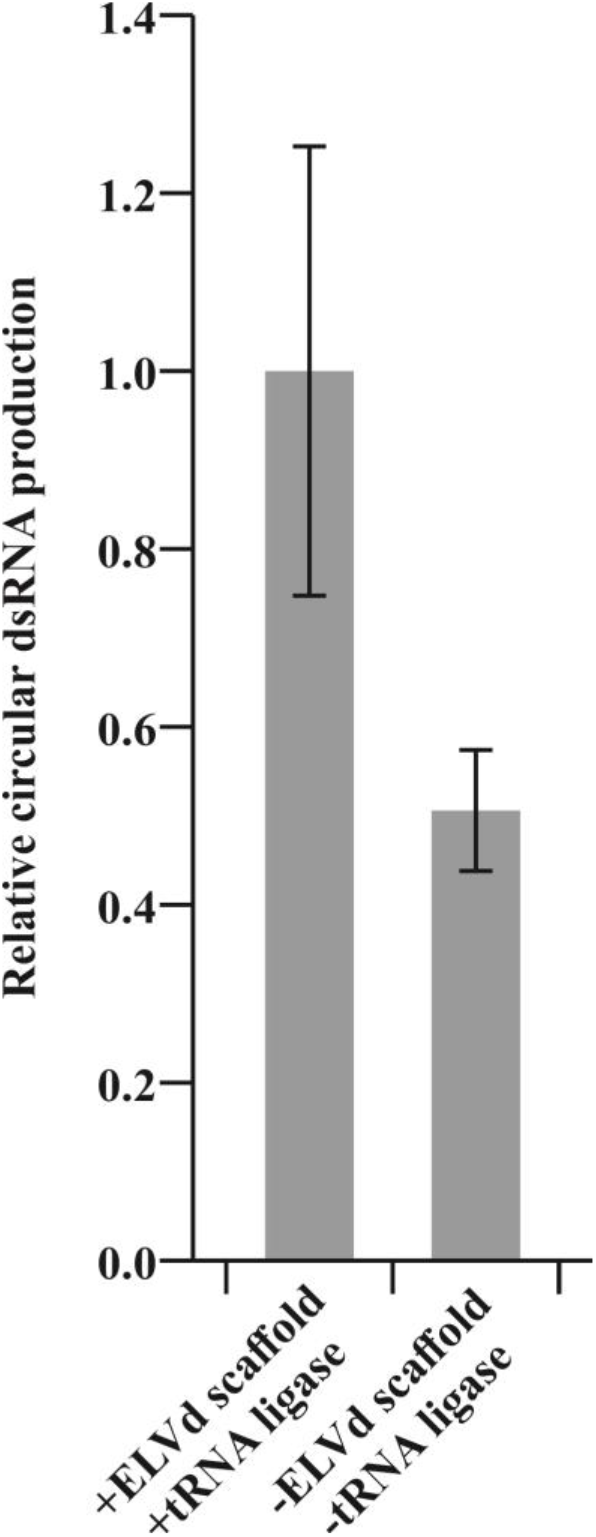
Effect of the the ELVd scaffold and the tRNA ligase on accumulation of a recombinant circular dsRNA. RNA preparations from *E. coli* clones cotransformed with p15LtRnlSm and a pLELVdPIE-derivative to produce a 100-bp dsRNA or the empty ligase plasmid (p15CAT) and a pLPIE-derivative (no ELVd scaffold) to produce the same 100 bp dsRNA were separated by denaturing PAGE. After staining the gels with ethidium bromide, recombinant circular dsRNA accumulation was quantified (in fluorescence arbitrary units) using an image analyzer. Normalized average fluorescence is plotted. Error bars represent standard deviation (n = 5).

**Supplemental Table S1.**
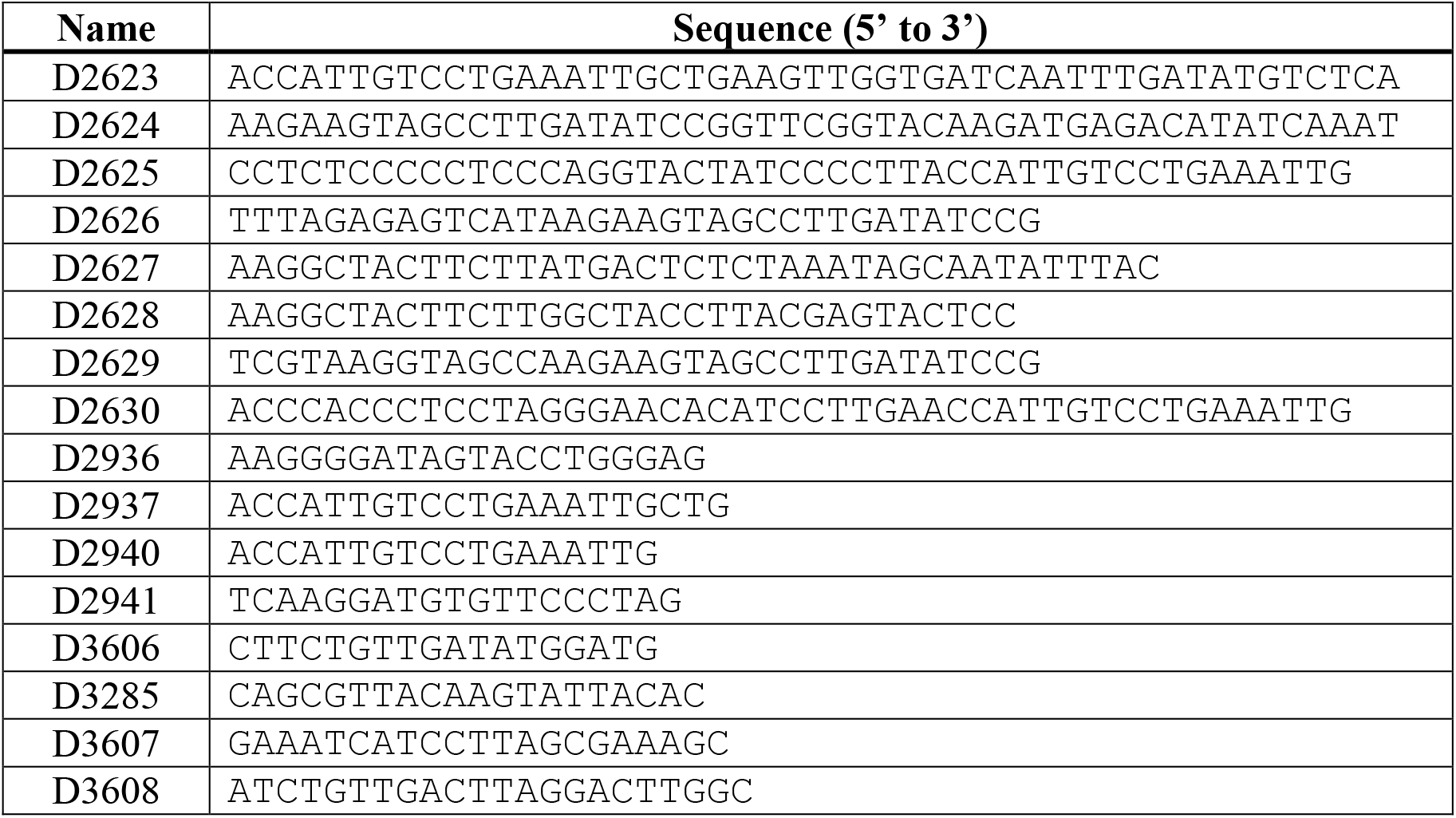
Primers used in the PCR amplifications to build expression plasmids pLELVd-DvSSJ1, pLELVdPIE-DvSSJ1 and pLPIE-DvSSJ1.

## Notes

### Competing Interest Statement

The authors have declared no competing interest.

